# BICC1 Interacts with PKD1 and PKD2 to Drive Cystogenesis in ADPKD

**DOI:** 10.1101/2024.08.27.608867

**Authors:** Uyen Tran, Andrew J Streets, Devon Smith, Eva Decker, Annemarie Kirschfink, Lahoucine Izem, Jessie M. Hassey, Briana Rutland, Manoj K Valluru, Jan Hinrich Bräsen, Elisabeth Ott, Daniel Epting, Tobias Eisenberger, Albert CM Ong, Carsten Bergmann, Oliver Wessely

## Abstract

Autosomal dominant polycystic kidney disease (ADPKD) is primarily of adult-onset and caused by pathogenic variants in *PKD1* or *PKD2*. Yet, disease expression is highly variable and includes very early-onset PKD presentations *in utero* or infancy. In animal models, the RNA-binding molecule Bicc1 has been shown to play a crucial role in the pathogenesis of PKD. To study the interaction between BICC1, PKD1 and PKD2 we combined biochemical approaches, knockout studies in mice and *Xenopus,* genetic engineered human kidney cells carrying BICC1 variants as well as genetic association studies in a large ADPKD cohort. We first demonstrated that BICC1 physically binds to the proteins Polycystin-1 and -2 encoded by *PKD1* and *PKD2* via distinct protein domains. Furthermore, PKD was aggravated in loss-of-function studies in *Xenopus* and mouse models resulting in more severe disease when *Bicc1* was depleted in conjunction with *Pkd1 or Pkd2*. Finally, in a large human patient cohort, we identified a sibling pair with a homozygous *BICC1* variant and patients with very early onset PKD (VEO-PKD) that exhibited compound heterozygosity of *BICC1* in conjunction with *PKD1 and PKD2* variants. Genome editing demonstrated that these *BICC1* variants were hypomorphic in nature and impacted disease-relevant signaling pathways. These findings support the hypothesis that BICC1 cooperates functionally with PKD1 and PKD2, and that *BICC1* variants may aggravate PKD severity highlighting RNA metabolism as an important new concept for disease modification in ADPKD.

## INTRODUCTION

Autosomal dominant polycystic kidney disease (ADPKD) is the most frequent life-threatening genetic disease and one of the most common Mendelian human disorders with an estimated prevalence of 1/400-1000.^1,2^ This equates to around 12.5 million affected individuals worldwide. About 5-10% of all patients requiring renal replacement therapy are affected by ADPKD. The majority of ADPKD patients carry a pathogenic germline variant in the *PKD1* or *PKD2* gene and present with the disease in adulthood.^2–4^ However, occasionally, ADPKD can manifest in infancy or early childhood [< 2 years, very-early onset ADPKD (VEO-ADPKD)], and in late childhood or early teenage years [2-16 years, early-onset ADPKD (EO-ADPKD)].^5,6^ VEO patients and fetuses often present with a Potter sequence and significant peri- or neonatal demise, which can be clinically indistinguishable from a typical Autosomal Recessive Polycystic Kidney Disease (ARPKD) presentation caused by *PKHD1* mutations.^7,8^ However, in contrast to VEO/EO-ADPKD, ARPKD kidneys invariably manifest as fusiform dilations of renal collecting ducts and distal tubules that usually remain in contact with the urinary system.^4^ Co-inheritance of an inactivating *PKD1* or *PKD2* mutation with an incompletely penetrant minor PKD allele *in trans* provides a likely explanation for VEO-ADPKD.^9^ In fact, we recently reported that the majority (70%) of VEO-ADPKD cases in an international diagnostic cohort had biallelic *PKD1* variants (i.e., a pathogenic variant *in trans* with a hypomorphic, low penetrance variant), while cases of biallelic *PKD2* and digenic *PKD1/PKD2* were rather rare.^10^ In line with the dosage theory for PKD,^11^ several other genes (*GANAB, DNAJB11, ALG8, ALG9, IFT140*) have been associated with a dominant, but late-onset atypical adult presentation and sometimes incomplete penetrance.^4,12–15^ However, not all VEO/EO-ADPKD patients can be explained by monogenic inheritance suggesting digenic or oligogenic inheritance causes.

Previous data from mouse, *Xenopus* and zebrafish suggest a crucial role for the RNA-binding protein Bicc1 in the pathogenesis of PKD, although *BICC1* mutations in human PKD have not been previously reported.^16–25^ BICC1 encodes an evolutionarily conserved protein that is characterized by 3 K-homology (KH) and 2 KH-like (KHL) RNA-binding domains at the N-terminus and a SAM domain at the C-terminus, which are separated by a by a disordered intervening sequence (IVS).^26–31^ The protein localizes to cytoplasmic foci involved in RNA metabolism and has been shown to regulate the expression of several genes such as *Pkd2, Adcyd6* and *Pkia* in the kidney.^22,32^ We now present data providing a mechanistic model linking BICC1 with the three major cystic proteins. We show that BICC1 physically interacts with the PKD1 (PC1) and the PKD2 (PC2) proteins in human kidney cells. We also demonstrate that *Pkd1* and *Pkd2* modifies the cystic phenotype in *Bicc1* mice in a dose-dependent manner and that Bicc1 functionally interacts with Pkd1, Pkd2 and Pkhd1 in the pronephros of *Xenopus* embryos. Finally, this interaction is supported by human patient data where *BICC1* alone or in conjunction with *PKD1* or *PKD2* is involved in VEO-PKD.

## METHODS

### Cell Culture and Biochemical Studies

The characterization of the interaction between BICC1, PC1 and PC2 as well as the analysis of the human *BICC1* variants were performed using standard approaches detailed in the Supplementary Methods.

### Animal Studies

Mouse and *Xenopus laevis* studies were approved by the Institutional Animal Care and Use Committee at the Cleveland Clinic Foundation and LSU Health Sciences Center (present and former employer of Dr. Wessely) and adhered to the National Institutes of Health Guide for the Care and Use of Laboratory Animals. Experimental design and data interpretation followed the ARRIVE1 reporting guidelines.^33^

### International Diagnostic Clinical Cohort

Research was performed following written informed consent and according to the declaration of Helsinki and oversight was provided by the Medizinische Genetik Mainz. DNA extraction was performed according to standard procedures (see Supplementary Methods for details).

### Statistical analysis

Data are presented as mean values ± SEM. Paired and unpaired two-sided Student’s t test or ANOVA were used for statistical analyses with a minimum of *P* < 0.05 indicating statistical significance.

Measurements were taken from distinct biological samples. Analyses were carried out using Prism 10 (Graphpad).

## RESULTS

### Interaction of BICC1 with PC1 and PC2

Loss of Pkd1 has been associated with lower Bicc1 expression in a murine model.^34^ Furthermore, Bicc1 has been shown to regulate *Pkd2* expression in cellular and animal models.^22,35,36^ However, whether this is due to direct protein-protein interactions between BICC1, PKD1 (PC1) and PKD2 protein (PC2) have not been investigated. In pilot experiments, BICC1 was detected by mass spectrometry in a pulldown assay from cells stably expressing a Polycystin-1 PLAT domain (Polycystin-1, Lipoxygenase, Alpha-Toxin)-YFP fusion.^37^ The direct binding between the PC1-PLAT domain and Bicc1 was confirmed using *in vitro* binding assays, but we also detected binding to the PC1 C-terminus (CT1) (**Supplementary Fig. S1a,c**).

Utilizing recombinant GST-tagged domains of PC1 and PC2, we demonstrated that Bicc1 binds to both proteins in GST-pulldown assays (**Fig. 1a, b**). In the case of PC1, myc-mBicc1 strongly interacted with its C-terminus (GST-CT1) but its interaction was abolished by a PC1-R4227X truncation mutation (GST-CT1-R4227X) (**Fig. 1b,c**). In the case of PC2, myc-mBicc1 associated with both recombinant GST N-terminal (GST-NT2) and C-terminal (GST-CT2) fusions. To investigate whether binding was direct or indirect, we performed *in vitro* binding assays using *in vitro* translated myc-Bicc1 and recombinant PC1 and PC2 domains. GST-pulldowns confirmed a direct interaction between myc-Bicc1 and GST-CT1 but not GST-CT1-R4227X (**Fig. 1d, e**). Similarly, myc-Bicc1 interacted directly with GST-NT2. While binding was stronger with the distal sequence (NT2 aa101-223) both N-terminal fragments contributed to the overall binding to Bicc1 (**Fig. 1d, e**). Interestingly, no direct interaction between Bicc1 and GST-CT2 was detected (**Supplementary Fig. S1b**) suggesting that the observed *in vivo* interaction with Bicc1 is indirect. Finally, immunoprecipitation using lysates from human kidney epithelial cells (UCL93) to assay endogenous, non-overexpressed proteins showed that PC1, PC2 and BICC1 form protein complexes *in vivo* (**Fig. 1f, g**).

**Figure 1.**
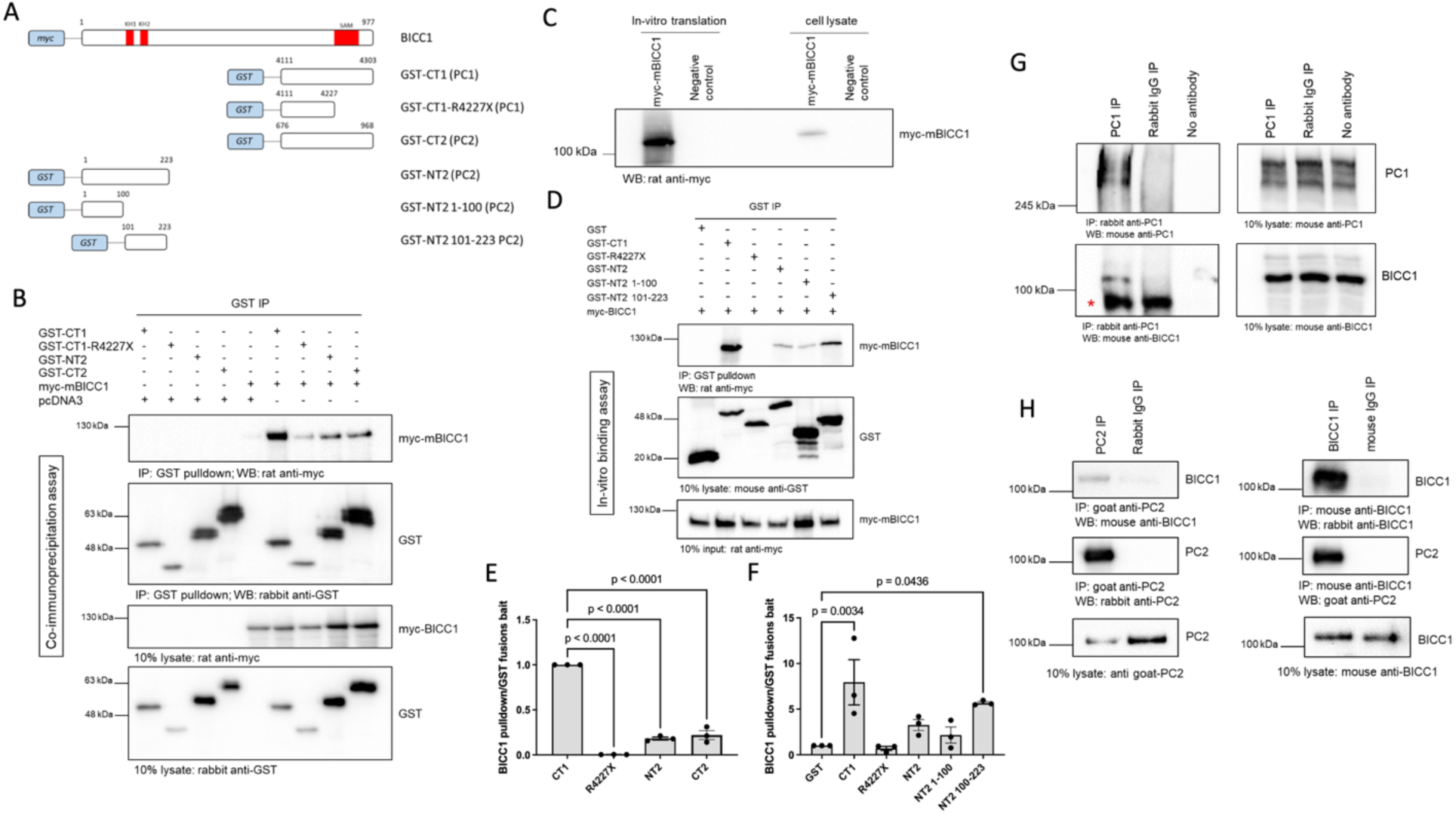
*Bicc1* Forms a Complex with Polycystin-1 and Polycystin-2. Full-length and deletion myc-tagged constructs of Bicc1 were co-expressed with either full-length HA-tagged PC1 or PC2 in HEK-293 cells and tested for their ability to interact by co-IP. (**a**) Schematic diagram of the constructs used in this experiment. (**b**) Western blot analysis following co-IP experiments, using GST tagged constructs as bait, identified protein interactions between PC1 or PC2 domains and *Bicc1*. pcDNA3 was included as a negative control. CT = C-terminus, NT = N-terminus, GST = Glutathione S-Transferase. Co-IP experiments (n=3) were quantified in (c). (**d**) Western blot analysis following *in-vitro* pulldown experiments, using purified GST tagged constructs as bait, identified direct protein interactions between PC1 or PC2 domains and *in vitro* translated myc-Bicc1. In-vitro binding experiments (n=3) were quantified in (e,f). (**g**) Western blot analysis following co-IP experiments, using a rabbit PC1 antibody (2b7) as bait, identified protein interactions between endogenous PC1 and BICC1 in UCL93 cells. A non-immune rabbit IgG antibody or no antibody was included as a negative control; * denotes a non-specific IgG band which is not present in the no antibody control lane. (**h**) Western blot analysis following co-IP experiments, using an anti-mouse Bicc1 or anti-goat PC2 antibody as bait, identified protein interactions between endogenous PC2 and BICC1 in UCL93 cells. Non-immune goat and mouse IgG was included as a negative control.

### Different Interaction Motifs for the Binding of Bicc1 to the Polycystins

To define the PC1/PC2 interaction domain(s) in Bicc1, we generated deletion constructs lacking the SAM domain (myc-mBicc1-ΔSAM, aa1-815) or the KH/KHL domains (myc-mBicc1-ΔKH, aa352-977) (**Fig. 2a**) and studied them by co-IP. Full-length PC1 co-immunoprecipitated with full-length myc-mBicc1 (**Fig. 2b,c**). Deleting the SAM domain did not significantly reduce the association to PC1 (∼55%, p=0.79) compared to full-length myc-mBicc1. However, an 8-fold stronger interaction was observed between full-length PC1 and myc-mBicc1-ΔKH compared to myc-mBicc1 or myc-mBicc1-ΔSAM. These results suggested that the interaction between PC1 and Bicc1 may involve the SAM but not the KH/KHL domains (nor the first 132 amino acids of Bicc1). Potentially the N-terminus (aa1-351) could have an inhibitory effect on PC1-BICC1 association.

**Figure 2.**
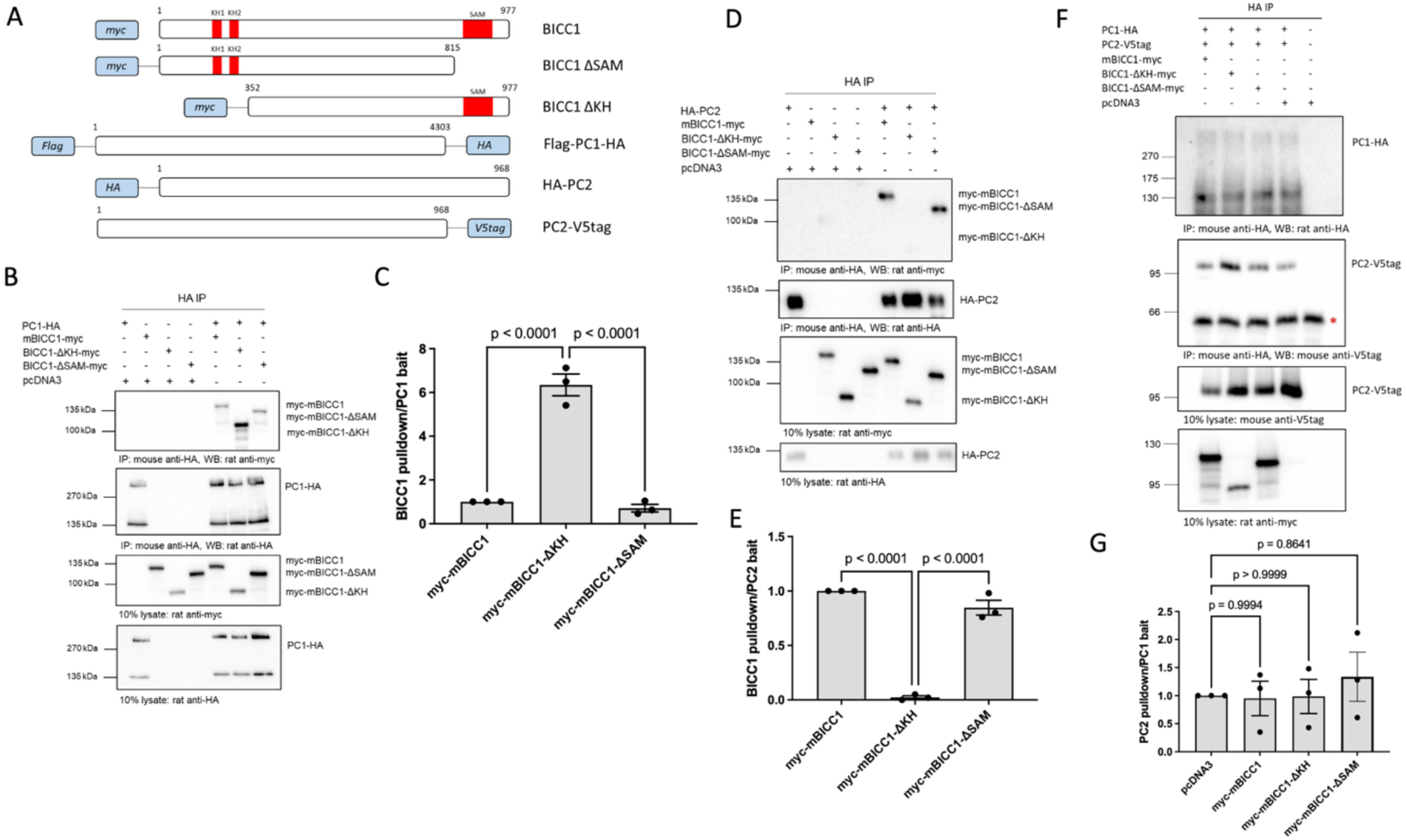
Interactions between *Bicc1* and Polycystin1/2 Require Different Binding Motifs. Full-length and deletion myc-tagged constructs of Bicc1 were co-expressed with either full-length HA-tagged PC1 or PC2 in HEK-293 cells and tested for their ability to interact by co-IP. (**a**) Schematic diagram of the constructs used in this set of experiments with the amino acid positions of full-length Bicc1 or the different deletions indicated. (**b,c**) Western blot analysis following co-IP experiments, using a PC1-HA-tagged construct as bait, identified protein interactions between PC1 and Bicc1 domains. pcDNA3 was included as a negative control (**b**). co-IP experiments (n=3) were quantified in (**c**). (**d,e**) Western blot analysis following co-IP experiments, using a PC2-HA tagged construct as bait, identified protein interactions between PC2 and Bicc1 domains (**d**). pcDNA3 was included as a negative control. Quantification of the co-IP experiments (n=3) is shown in (**e**). (f, g) Western blot analysis following co-IP experiments, using a PC1-HA-tagged construct as bait. The interaction between PC1 and PC2 was not altered in the presence of either full-length Bicc1 or its deletion domains. pcDNA3 was included as a negative control. Asterix represents non-specific interaction with mouse IgG. **(f)**. co-IP experiments (n=3) were quantified in **(g).** One way ANOVA comparisons were performed to assess significance and *P* values are indicated. Error bars represent standard error of the mean.

Similar experiments were performed to define the Bicc1 interacting domains for PC2 (**Fig. 2d,e**). Full-length PC2 (PC2-HA) interacted with full-length myc-mBicc1. Unlike PC1, PC2 interacted with myc-mBicc1-ΔSAM, but not myc-mBicc1-ΔKH suggesting that PC2 binding is dependent on the N-terminal domains (aa1-351) but not the SAM domain or distal C-terminus (aa816-977). Co-expression of Bicc1 deletion constructs lacking the SAM domain (myc-mBicc1-ΔSAM) or the KH domains (myc-mBicc1-ΔKH) however had no effect on the interaction of PC1 with PC2 in co-immunoprecipitation assays (**Fig 2f,g**) suggesting that these interactions are not mutually exclusive.

### Cooperativity of BICC1 with other PKD genes

Since our biochemical analysis indicated a direct interaction between BICC1, PC1 and PC2, we wondered whether this is biologically relevant. If this were the case, BICC1 should cooperate with other PKD genes and reducing BICC1 activity in conjunction with reducing either PKD1 or PKD2 activity should still cause a cystic phenotype. We first addressed this question in the *Xenopus* system (**Fig. 3**), which is an easily manipulatable model to study PKD. The PKD phenotype in frog is characterized by dilated kidney tubules, the loss of the expression of the sodium bicarbonate cotransporter 1 (Nbc1) in the in the distal tubule and the emergence of body-wide edema as a sign of a malfunctioning kidney.^21,22,38,39^ Knockdown of Bicc1, Pkd1, Pkd2 or the ARPKD gene Pkhd1 caused a PKD phenotype (**Fig. 3e-i”** and **Supplementary Fig. S2a**). The latter, *Pkhd1* was included to assay not only ADPKD, but also ARPKD, which is generally thought to disturb the same cellular mechanisms. To test whether Bicc1 cooperated with the PKD genes we then performed combined knockdowns. We titrated each of the four MOs to a concentration that on its own resulted in little phenotypic changes upon injection into *Xenopus* embryos (**Fig. 3j,k** and **Supplementary Fig. S2b**). However, combining *Bicc1-MO1+2* with *Pkd1-sMO*, *Pkd2-MO* or *Pkhd1-sMO* at suboptimal concentrations resulted in the re-emergence of a strong PKD phenotype.

**Figure 3.**
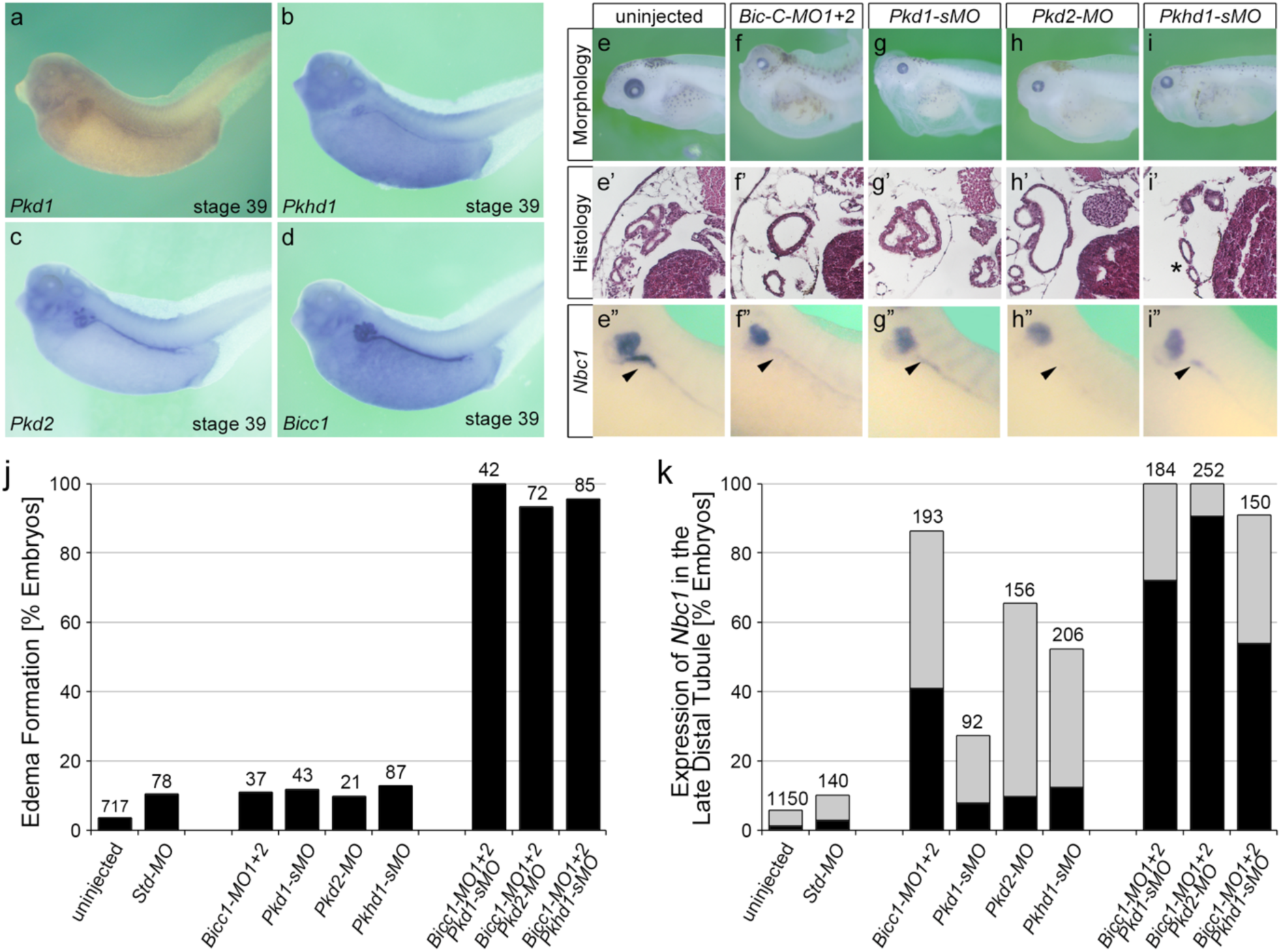
Cooperativity of Bicc1 and PKD Genes in *Xenopus*. (**a-d**) mRNA expression of *Pkd1*, *Pkhd1, Pkd2* and *Bicc1* in the *Xenopus* pronephros at stage 39. (**e-i”**) Knockdown of Bicc1 (**f-f”**), Pkd1 (**g-g”**), Pkd2 (**h-h”**) and Pkhd1 (**i-i”**) by antisense morpholino oligomers (MOs) results in a PKD phenotype compared to uninjected control *Xenopus* embryos (**e-e”**). The phenotype is characterized by the formation of edema due to kidney dysfunction (**e,f,g,h,i**; stage 43), the development of dilated renal tubules (**e’,f’,g’,h’,i’**; stage 43) and the loss of *Nbc1* in the late distal tubule by whole mount *in situ* hybridizations (arrowheads in **e”,f”,g”,h”,i”**; stage 39). **(j,k)** To examine cooperativity, *Xenopus* embryos were injected with suboptimal amounts of the MOs, either alone or in combination, and analyzed for edema formation at stage 43 (**j**) and the expression of *Nbc1* at stage 39 (**k**) with gray bars showing reduced and black bars showing absent *Nbc1* expression in the late distal tubule. Data are the accumulation of multiple independent fertilizations with the number of embryos analyzed indicated above each condition.

While injections with individual MOs developed edema in about 10% of the embryos, co-injections caused edema formation in almost 100% of the embryos (**Fig. 3j**, last 3 columns). A similar result was seen for the expression of *Nbc1* in the late distal tubule, where individual MO injections showed some changes in gene expression, but double MO injections had a highly synergistic effect resulting in a near complete loss of *Nbc1* (**Fig. 3k**). Of note, this approach is based on a sensitized biological readout, but not on reducing expression levels to a fixed amount (e.g., 50%).

We next investigated whether a similar cooperation between Bicc1 and Pkd1 or Pkd2 can be observed in genetic mouse models. We initially focused on Bicc1 and Pkd2. Both *Bicc1* and *Pkd2* knockout mice develop cystic kidneys as early as E15.5.^22,40^ As this is the earliest time point cystic kidneys can be observed, crossing those strains did not allow us to assess cooperativity (data not shown). Moreover, like in the case of compound *Pkd1/Pkd2* mutants,^41^ kidneys from *Bicc1^+/-^:Pkd2^+/-^* did not exhibit cysts (data not shown). Thus, we instead used mice carrying the Bicc1 hypomorphic allele *Bpk*, which develop a cystic kidney phenotype postnatally.^18,42^ To assess cooperativity, we removed one copy of *Pkd2* in the *Bpk* mice. Comparing the kidneys of *Bicc1^Bpk/Bpk^:Pkd2^+/-^*to those of *Bicc1^Bpk/Bpk^:Pkd2^+/+^*at postnatal day P14 revealed that the compound mutant kidneys were larger and more translucent (**Fig. 4a**) and the kidney/body weight ratios (KW/BW) were significantly increased (**Fig. 4b**). Moreover, analyzing survival the compound mutants showed a trend towards an earlier demise (**Supplementary Table S1**). We did not detect sex differences in the phenotype (**Supplementary Figure S3c**). Yet, the reduction in *Pkd2* gene dose affected the progression of the disease, but not its onset. Performing the same analysis at postnatal day P4 did not show any differences (**Fig. 4c**).

**Figure 4:**
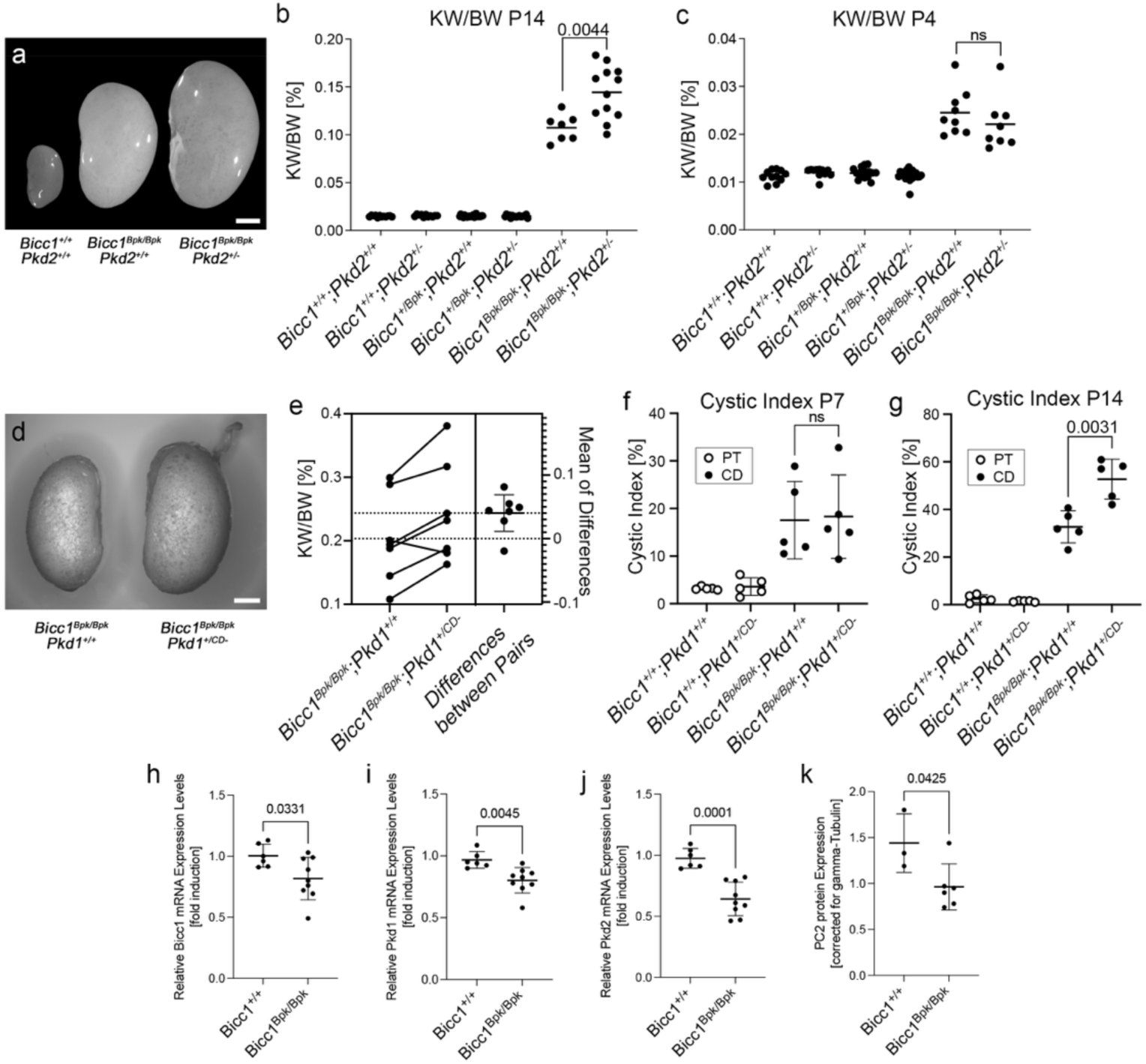
Cooperativity of Bicc1 and Pkd1 and Pkd2 in Mouse. (**a-c**) Bicc1 and Pkd2 interact genetically. Offspring from *Bicc1;Pkd2* compound mice at postnatal day P4 and P14 are compared by outside kidney morphology at postnatal day P14 (**a,** scale bar is 2 mm), and kidney to body weight ratio (KW/BW) at P14 (**b**) and P4 (**c**). (**d-g**) Bicc1 and Pkd1 interact genetically. *Bicc1;Pkd1* compound mice are compared by outside kidney morphology at P14 showing a kidney from *Bicc1^Bpk/Bpk^:Pkd1^+/+^*and a *Bicc1^Bpk/Bpk^:Pkd1^+/CD-^*littermates (**d,** scale bar is 2 mm, no kidney of a wildtype littermate is shown, as it was not present in the litter), estimation plot of KW/BW ratio comparing littermates at P14 with a *P*-value = 0.092 (**e**), and cystic index, i.e. percent of of proximal tubules (PT) and collecting ducts (CD) cysts in respect to the total kidney area at P7 (**f**) and P14 (**g**). Two-sided paired t-tests were performed to assess significance, and the *P*-values are indicated; error bars represent standard deviation. (**h-k**) qRT-PCR analysis for *Bicc1*, *Pkd1*, and *Pkd2* expression (**h-j**) and quantification of the PC2 expression levels by Western blot (**k**) in kidneys at P4 before the onset of a strong cystic kidney phenotype. Data were analyzed by t-test and the *P*-values are indicated. Please note that the y-axes of the different panels are intentionally different to best visualize the changes between the groups analyzed.

Next, we performed a similar mouse study for *Pkd1* using the *Pkd1^Fl/Fl^:Pkhd1-Cre* line described previously^43^ (in the following referred to as *Pkd1^CD^*^-^). This mouse line eliminates *Pkd1* postnatally in the collecting ducts. Similar to the *Bicc1/Pkd2* scenario, when removing one copy of *Pkd1* in the collecting ducts, the *Bicc1^Bpk/Bpk^:Pkd1^+/CD-^*appeared larger when comparing kidneys from littermates (**Fig. 4d**) and littermates exhibited statistically significant differences in KW/BW ratio (**Fig. 4e**). Yet, the phenotype was rather subtle and aggregating all the data did not show differences in KW/BW ratios between *Bicc1^Bpk/Bpk^:Pkd1^+/+^*and *Bicc1^Bpk/Bpk^:Pkd1^+/CD-^*mice (**Supplementary Fig. S3d**). Thus, to further corroborate the genetic interaction, we determined the cystic index for proximal tubules and collecting ducts using LTA and DBA staining, respectively. This showed an increase in collecting duct cysts upon removal of one copy of *Pkd1* (**Fig. 4g**). Like in the case of *Pkd2*, the phenotype seems to be correlated with cyst expansion and not the onset, as there was no difference at postnatal day P7 (**Fig. 4f**) and we did not detect increased mortality in the compound mutants (**Supplementary Table S2)**. It is noteworthy that neither the *Bicc1/Pkd2* nor the *Bicc1/Pkd1* compound mutants showed an aggravated kidney function based on Blood Urea Nitrogen (BUN) levels (**Supplementary Fig. S3a,b,e**) likely due to the aggressive nature of the *Bicc1^Bpk/Bpk^* phenotype. Of note, due to the different genetic approaches using a *Pkd2* null allele and a conditional *Pkd1* allele, the outcomes of the two crosses cannot be directly compared. Yet, these *in vivo* data support our biochemical interaction data and demonstrated that *Bicc1* cooperates with *Pkd1* and *Pkd2*.

Finally, to better understand how Bicc1 would exert such a phenotype, we analyzed the expression of the PKD genes in the *Bicc1^Bpk/Bpk^*mice. We have previously demonstrated that *Pkd2* levels are reduced in a complete Bicc1 null mice.^22^ Performing qRT-PCR of kidneys from wildtype and *Bicc1^Bpk/Bpk^* at P4 (i.e. before the onset of a strong cystic phenotype), revealed that *Bicc1, Pkd1* and *Pkd2* were statistically significantly down-regulated (**Fig. 4h-j**). The effect on *Pkd2* mRNA was confirmed by protein analysis for PC2 (**Fig. 4k** and **Supplementary Fig. S3f**).

### *BICC1* Variants in Patients with early and severe Polycystic Kidney Disease

To evaluate whether these interactions are relevant for human PKD, we analyzed an international cohort of 2,914 PKD patients by massive parallel sequencing (MPS) ^44,45^ focusing on VEO-ADPKD patients with the hypothesis that *BICC1* variants may lead to a more severe and earlier PKD phenotype. While variants in *BICC1* are very rare, we could identify two patients with *BICC1* variants harboring an additional *PKD2* or *PKD1* variant *in trans*, respectively. None of these *BICC1* variants were detected in the control cohort or in non-VEO-ADPKD patients. Moreover, besides the variants reported below, the patients had no other variants in any of other PKD genes or genes which phenocopy PKD including *PKD1*, *PKD2*, *PKHD1*, *HNF1ß*, *GANAB*, *IFT140*, *DZIP1L*, *CYS1*, *DNAJB11*, *ALG5*, *ALG8*, *ALG9*, *LRP5*, *NEK8*, *OFD1* or *PMM2*.

The first patient was severely and prenatally affected demonstrating a Potter sequence with huge echogenic kidneys and oligo-/anhydramnios. Autopsy confirmed VEO-ADPKD with absence of ductal plate malformation invariably seen in ARPKD. The fetus carried the *BICC1* variant (c.2462G>A, p.Gly821Glu) inherited from his father, who presented with two small renal cysts in one of his kidneys, and a *PKD2* variant (c.1894T>C, p.Cys632Arg) that arose *de novo* (**Fig. 5a**). Individual *in silico* predictions (SIFT, Polyphen2, CADD, Eigen-PC, FATHMM, GERP++ RS and EVE), meta scores (REVEL, MetaSVM, and MetaLR) and other protein function predictions (PrimateAI, AlphaMissense, ESM1b and ProtVar) indicate that this *PKD2* missense variant is likely pathogenic (**Supplementary Table S3**). Moreover, structural analysis suggests that the hydrophilic substitution may interfere with the Helix S5 pore domain of PKD2 and change its ion channel function (**Fig. 5b,c**). Finally, *PKD2* p.Cys632Arg has been previously reported as part of a PKD2 pedigree and implicated as a critical determinant for Polycystin-2 function.^46,47^ On the other hand, the *BICC1* p.Gly821Glu variant is located in an intrinsically disordered domain of BICC1 between the KH and the SAM domains (**Fig. 6f**). To address whether the variant is hypomorphic, we used CRISPR-Cas9-mediated gene editing to generate HEK293T cells lacking BICC1 or harboring the p.Gly821Glu mutation (BICC1-G821E). These cells were analyzed for their impact on the translation of *PKD2*, a well-established target of Bicc1.^22^ As shown in **Fig. 5d,e** PC2 protein levels were strongly reduced in two independent HEK293T BICC1-G821E cells when compared to unedited HEK293T cells. Most notably, the PC2 levels were comparable to the levels found in HEK293T carrying a *BICC1* null allele (HEK293T BICC1-KO) (**Supplementary Figure S2c,d**). Based on these data we hypothesize that the major disease effect results from the pathogenic *PKD2* variant but is aggravated by the *BICC1* variant.

**Figure 5.**
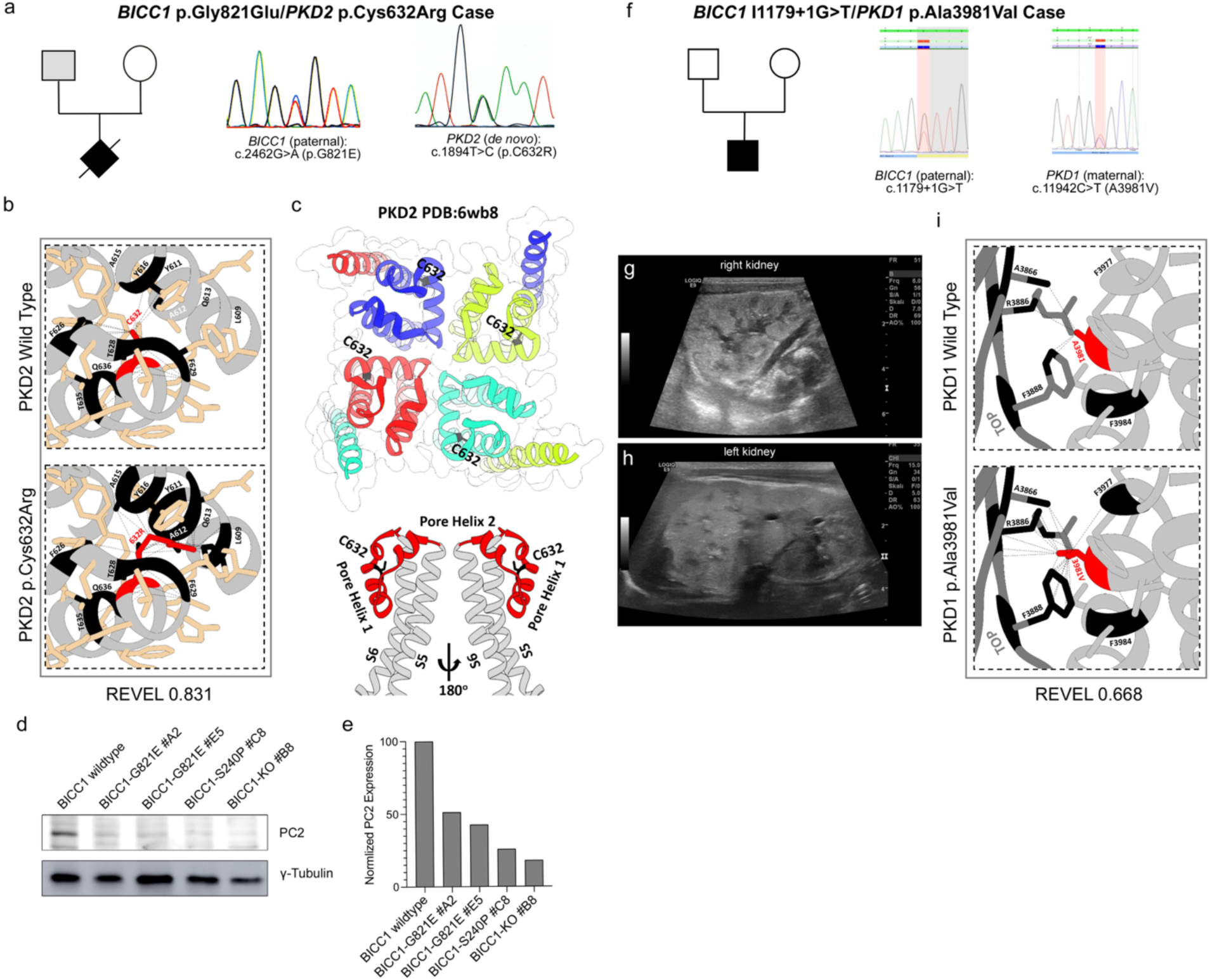
Identification of Human *BICC1* Variants. (**a-c**) *BICC1* p.G821E/*PKD2* p.C632R case with pedigree and the electropherograms (**a**), the structural analysis of the PKD2 showing the local structure around the cysteine at position 632 (indicated in red) and its putative impact in the variant including the REVEL score (**b**) as well as its location within the global PC2 structure highlighting the potential of the variant impacting the PC2 ion channel function (**c**). (**d,e**) Western blot analysis for PC2 comparing wildtype HEK293T, HEK293T BICC1 p.Gly821Glu (BICC1-G821E), HEK293T BICC1 p.Ser240Pro (BICC1-S240P) and HEK293T BICC1 knockout (BICC1-KO) cells and quantification thereof. ◻-Tubulin was used as loading control. (**f-i**) *BICC1* I1179+1G>T/*PKD1* p.Ala3981Val case with pedigree and the electropherograms (**f**), the ultrasound analysis of the left and right kidneys (**g,h**) and the structural analysis of the PC1 showing the local structure around the alanine at position 3981 (indicated in red) and its putative impact in the variant including the REVEL score (**i**).

**Figure 6:**
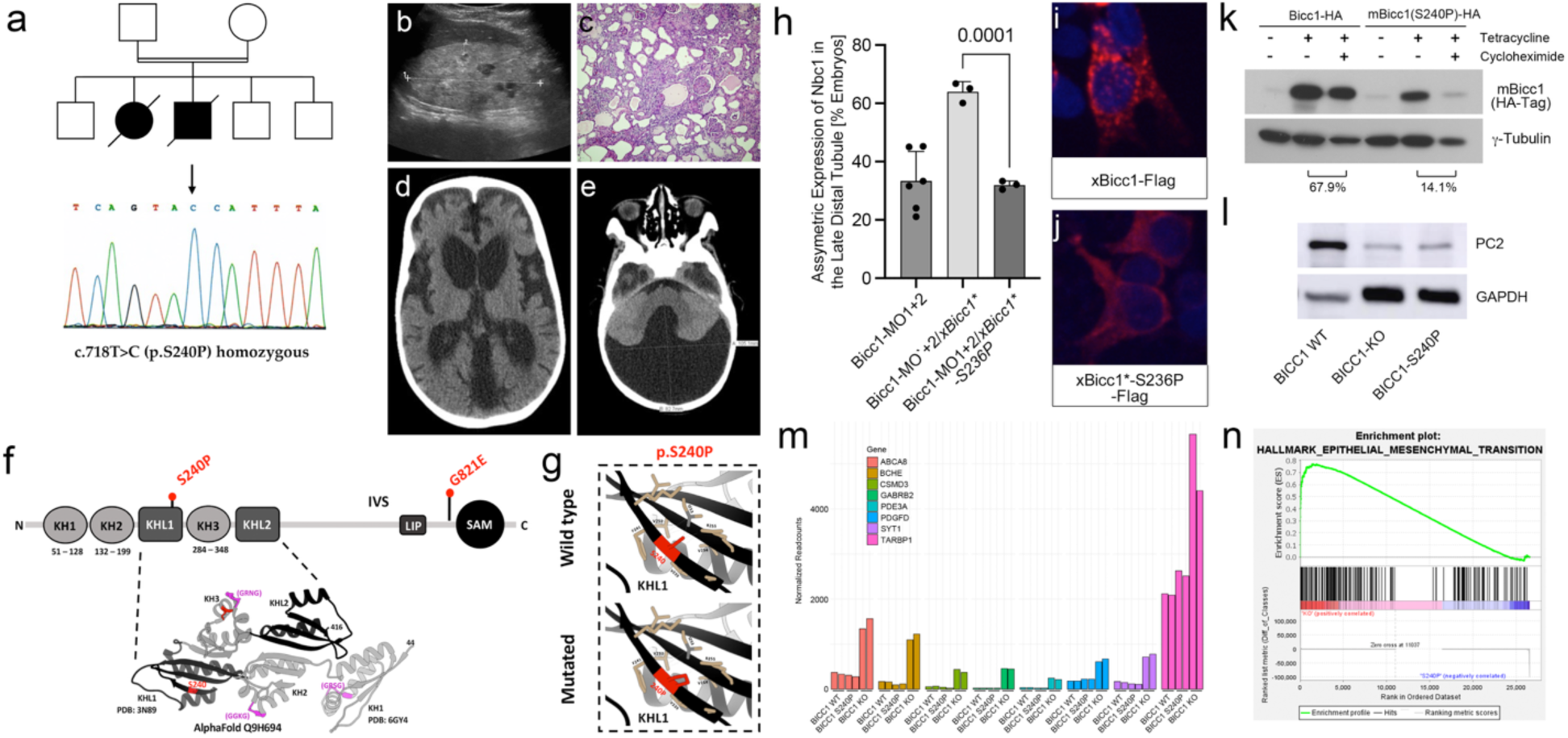
The Homozygous *BICC1* p.Ser240Pro Variant is a Hypomorphic Cystic Disease-Causing Variant. (**a-e**) Consanguineous multiplex pedigree with two siblings affected by VEO-ADPKD identified the homozygous *BICC1* missense variant c.718T>C (*BICC1* p.Ser240Pro) absent from gnomAD and other internal and public databases. Electropherogram is shown in (**a**). The affected girl presented at a few months of age with renal failure and enlarged polycystic kidneys that lacked corticomedullary differentiation (**c**). Histology after bilateral nephrectomy showed polycystic kidneys more suggestive of ADPKD than ARPKD without any dysplastic element. Her younger brother exhibited enlarged hyperechogenic polycystic kidneys prenatally by ultrasound (**b**). In addition, in his early infancy arterial hypertension and a Dandy-Walker malformation with a low-pressure communicating hydrocephalus were noted (**d,e**). (**f**) Ribbon diagram and schematic diagram of BICC1 showing the KH, KHL and SAM domains. The two *BICC1* variants identified in this study, BICC1 p.Ser240Pro (S240P) and BICC1 p.Gly821Glu (G821E) are indicated in red. (**g**) Solid boxes correspond to local impacts of p.Ser240Pro (p.S240P) on BICC1 structure, interactions are labeled as dashed lines (pseudobonds). GXXG motifs colored in magenta, representative missense variant residues colored in red and residues adjacent to selected variant (<5Å) colored in tan. (**h**) Rescue experiments of *Xenopus* embryos lacking BicC1 by co-injections with the wild type or mutant constructs. Embryos were scored for the re-expression of *Nbc1* in the late distal tubule by whole mount *in situ* hybridizations. Quantification of at least 3 independent experiments is shown. **(i,j**) HEK293T cells were transfected with Flag-tagged constructs of wild type or mutant Bicc1 and the subcellular localization of Bicc1 was visualized (red). Nuclei were counterstained with DAPI (blue). (**k**) Protein stability analysis using tetracycline-inducible HEK293T cells comparing the expression levels of Bicc1 and Bicc1-S240P 24 hours after removal of tetracycline and addition of cycloheximide. ◻-Tubulin was used as loading control. The percentage of protein destabilization because of protein synthesis inhibition by cycloheximide is indicated. (**l**) Western Blot analysis of wildtype HEK293T, cells lacking BICC1 (BICC1-KO) and isogenic cells with the BICC1 p.Ser240Pro (BICC1-S240P) variant for PC2 expression. GAPDH was used as loading control. (**m,n**) Bar graph of the mRNA-seq transcriptomic analysis comparing BICC1 wildtype, knockout and S240P isogenic HEK293T cells showing the eight most significantly upregulated transcripts (based on their Padj levels) in the BICC1 KO cells (**m**). For each gene, the normalized expression levels from each of the 6 samples (2 wildtype, KO and 240P each) is shown. (**n**) GSEA plot showing the enrichment of the Hallmark Epithelial_Mesenchymal_Transition data set in the BICC1-KO cells *vs.* the BICC1-S240P cells.

The second patient presented perinatally with massively enlarged hyperechogenic kidneys, while the parents, both in their thirties, and the remaining family members were reported to be healthy (**Fig. 5f-h**). He carried a paternal canonic *BICC1* pathogenic splicing variant (c.1179+1G>T), which is likely pathogenic as the protein is truncated after exon 10, and a novel heterozygous *PKD1* variant (c.11942C>T, p.Ala3981Val) which has not been previously reported (**Fig. 5f**). While the *PKD1* variant appears minor in its amino acid change (i.e. Ala to Val), *in silico* analyses using individual predictions (SIFT, Polyphen2, CADD and EVE), Meta scores (REVEL) and other protein function predictions (PrimateAI, Alphamissense and ESM1b) indicate that the missense variant is likely pathogenic (**Supplementary Table S3**). Structural analyses suggest that although the Ala3981Val variant does not destabilize the Helix structure, its contact with the TOP domain could interfere with domain flexibility and PC1 complex assembly.

### A Sibling Pair of PKD Patients with a Homozygous *BICC1* Variant

The most insightful finding for a critical role for BICC1 in human PKD was the discovery of a homozygous *BICC1* variant in a consanguinous Arab multiplex pedigree, two siblings, a boy and a girl, diagnosed with VEO-ADPKD (**Fig. 6a-e**). The affected female presented at a few months of age with kidney failure and enlarged polycystic kidneys that lacked corticomedullary differentiation. Histology after bilateral nephrectomy showed polycystic kidneys more suggestive of ADPKD than ARPKD without any dysplastic element (**Fig. 6c**). Her younger brother exhibited enlarged hyperechogenic polycystic kidneys antenatally by ultrasound (**Fig. 6b**). In addition, during early infancy, arterial hypertension and a Dandy-Walker malformation with a low-pressure communicating hydrocephalus were noted (**Fig. 6d,e**). By customized MPS, we identified the homozygous missense *BICC1* variant (c.718T>C, p.Ser240Pro) (**Fig. 6a**). This variant was absent from gnomAD and fully segregated with the cystic phenotype present in this family. It results in a non-conservative change from the aliphatic, polar-hydrophilic serine to the cyclic, apolar-hydrophobic proline located in the second beta sheet of the first KHL1 domain and very likely disrupts the beta sheet and thus the RNA-binding activity of Bicc1 (**Fig. 6f,g** and **Supplementary Table S4**). In the more severely affected younger brother, we also detected an additional heterozygous *PKD2* variant (c.1445T>G, p.Phe482Cys), which results in a non-conservative change from phenylalanine to cysteine (**Supplementary Table S3)**. It was previously reported that this PC2 Phe482Cys variant exhibited altered kinetic PC2 channel properties, increased expression in IMCD cells and a different subcellular distribution when compared to wild-type PC2,^48^ These features suggested altered properties of this PC2 variant, yet its contribution to the case reported here remain untested.

Unfortunately, both siblings passed away and besides DNA and the phenotypic analysis described above neither human tissue nor primary patient-derived cells could be collected. Thus, to validate the pathogenicity of this point mutation, we turned to the amphibian model of PKD.^21,22^ In *Xenopus*, knockdown of Bicc1 using antisense morpholino oligomers (*Bicc1-MO1+2*) causes a PKD phenotype, which can be rescued by co-injection of synthetic mRNA encoding *Bicc1*.^21^ To test whether *BICC1* p.Ser240Pro had lost its biological activity, we introduced the same mutation into the *Xenopus* gene where the Ser is located at position 236 of the *Xenopus* gene (in the following referred to as *xBicC1*-S236P*). *Xenopus* embryos were injected with *Bicc1-MO1+2* at the 2-4 cell stage followed by a single injection of 2 ng wild type or *xBicc1*-S236P* mRNAs at the 8-cell stage. At stage 39 (when kidney development has been completed) embryos were analyzed by whole mount *in situ* hybridization for the expression of *Nbc1* in the late distal tubule of the pronephric kidney, one of the most reliable readouts for the amphibian PKD phenotype.^21^ As shown in **Fig. 6h**, wild type *Bicc1* mRNA restored expression of *Nbc1* on the injected side in 63% of the embryos. However, *xBicc1*-S236P* did not have any effect, and the embryos were indistinguishable from those injected with the *Bicc1-MO1+2* alone. This suggested that *xBicc1*-S236P* was functionally impaired. To address this hypothesis, we first assessed the subcellular localization of Bicc1 to foci that are thought to be involved in mRNA processing.^19,22,30,49^ Transfection of Flag-tagged Bicc1 (*xBicc1*-S236P-Flag*) into HEK293T cells reproduced this pattern (**Fig. 6i**). Surprisingly, xBicc1*-S236P-Flag was no longer detected in these cytoplasmic foci but rather homogenously dispersed throughout the cytoplasm (**Fig. 6j**). Western blot analysis demonstrated that this was accompanied by a reduction in protein levels (**Fig. 6k**). *In vitro* transcription/translation detected no differences between the proteins suggesting that the wildtype and Bicc1 S236P-Flag are translated equivalently (data not shown). Yet, in an *in vivo* pulse-chase experiments a mBICC1 p.Ser240Pro variant was less stable than its wildtype counterpart (**Fig. 6k**). However, whether the reduced protein level was due to an inherent instability of the mutant protein or a consequence of its mislocalization remains to be resolved. Finally, as in the case of BICC1 p.Gly821Glu we engineered HEK293T cells to harbor the *BICC1* p.Ser240Pro variant (BICC1-S240P). Western blot analysis demonstrated a reduction in PC2 levels in the BICC1-S240P cells when compared to unedited cells and that this reduction was comparable to PC2 levels in BICC1-KO cells (**Figs. 5d,e** and **6l**).

Finally, to determine to what extent the *BICC1* p.Ser240Pro variant differs from a *BICC1* loss of function allele, we performed mRNA sequencing (mRNA-seq) of the genetically engineered HEK293T cells. Differential gene expression analysis identified several genes that were differentially up- or down-regulated in the BICC1-S240P and the BICC1-KO cells compared to their unedited counterpart (**Supplementary Fig. 4a,e**). Approximately 24% and 18% of the differentially expressed genes were shared between BICC1-S240P or the BICC1-KO cells, respectively (**Supplementary Fig. S4b,f**). Yet, a substantial number of genes were specific to either cell line. The BICC1-S240P-enriched/depleted transcripts were generally also enriched/depleted in the BICC1-KO cells but did not reach statistical significance (**Supplementary Fig. 4c,g**). Conversely, many of the BICC1-KO enriched transcripts were specifically enriched/depleted in the BICC1-KO cells and not in the BICC1-S240P cells (**Fig. 6m** and **Supplementary Fig. 4d**). This suggested that there are qualitative differences between a null phenotype and the *BICC1* p.Ser240Pro variant, supporting our hypothesis that *BICC1* p.Ser240Pro acts as a hypomorph. Indeed, Gene Set Enrichment Analysis (GSEA) using the hallmark gene sets and comparing BICC1-KO and BICC1-S240P cells revealed a statistically significant enrichment for the Hallmark_Epithelial_Mesenchymal_Transition set (**Fig. 6n** and **Supplementary Table S5**), a pathway previously implicated in ADPKD.^50,51^

## DISCUSSION

BICC1 has been extensively studied in multiple animal models, which have suggested a critical role for BICC1 in several different developmental processes and in tissue homeostasis.^26^ This study functionally implicates it to human disease in general and PKD in particular by identifying the homozygous *BICC1* p.Ser240Pro variant, which was sufficient to cause a cystic phenotype in a sibling pair of human PKD patients. It is noteworthy that another study identified heterozygous *BICC1* variants in two patients with mildly cystic dysplastic kidneys.^23^ Yet, both variants were also present in one of the unaffected parents. While such a situation is extremely rare and does not significantly contribute to the mutational load in ADPKD or ARPKD, it demonstrated that loss of BICC1 is sufficient to cause PKD in humans. In addition, variants in *BICC1* and *PKD1 and PKD2* co-segregated in PKD patients from an International Clinical Diagnostic Cohort. While we have not yet shown the impact of each variant when introduced in a compound heterozygous situation, we postulate that PKD alleles *in trans* and/or *de novo* exert an aggravating effect and contribute to polycystic kidney disease. A reduced dosage of PKD proteins would severely disturb the homeostasis and network integrity, and by this correlates with disease severity in PKD. ADPKD is quite heterogeneous and - even within the same family - shows quite some phenotypic variation.^52,53^ It is thought that stochastic inputs, environmental factors and genetics influence PKD.^53^ The demonstrated interaction of BICC1, PC1 and PC2 now provides a molecular mechanism that can explain some of the phenotypic variability in these families. Of note, while our mouse studies support cooperation between Bicc1, Pkd1 and Pkd2, genetic proof for Bicc1 acting as a disease modifier, i.e. reduction of Bicc1 activity in a homozygous *Pkd1* or *Pkd2* background remains outstanding.

The second important aspect of this study is that BICC1 emerges as a central in the regulation of PKD1/PKD2 activity. Functional studies reported here and previously^22,35,36^ demonstrate that Bicc1 regulates the expression of *Pkd1* and *Pkd2*. Moreover, we now show that Bicc1 and PC1/PC2 physically interact and that lowering the expression levels of both proteins is sufficient to cause a PKD phenotype in frogs. Finally, the reduction of the gene dose for Pkd1 or Pkd2 in a hypomorphic mouse allele of *Bicc1* results in a more severe cystic kidney phenotype. These results in the kidney are paralleled and augmented in studies of left/right patterning, where Pkd2 can activate Bicc1 and where Bicc1 triggers critical aspects in establishing laterality.^19,30,54,55^ Thus, it is tempting to speculate that BICC1/PKD1/PKD2 are components of a critical regulatory network in maintaining epithelial homeostasis.

BICC1 has emerged as an important posttranscriptional regulator modifying gene expression through modulating the effects of microRNAs (miRNAs), regulating mRNA polyadenylation and translational repression and activation.^22,26,32,56–59^. While *PKD2* is the most appealing target in respect to ADPKD,^22^ there are undoubtable others (e.g. adenylate cyclase-6)^32^ that may be equally critical. Lastly, Bicc1 has been implicated in the regulation of miRNAs such as those of the *miR-17* family.^22^ This is of particular interest as a connection between *miR-17* activity and PKD is well-established.^60–66^ Both *Pkd1* and *Pkd2* mRNA are targeted by *miR-17*,^67^ and an *anti-miR-17* oligonucleotide is being developed as a PKD therapeutic.^68^ While we have shown that Bicc1 and *miR-17* targets *Pkd2* mRNA,^22^ a similar scenario for *Pkd1* is possible, but not yet shown. Thus, a tempting hypothesis is that the interaction between Bicc1, Pkd1 and Pkd2 and miRNAs - even though not examined in this study – compartmentalizes Bicc1’s activity where Bicc1 is posttranscriptionally inactive when complexed to Pkd1/Pkd2, but modulates Pkd1/Pkd2 expression when unbound. Such a regulatory complex could be responsible for several of the aspects of human ADPKD. In the future, it would be interesting to see how Bicc1 and its posttranscriptional targets are integrated and together contribute towards preventing kidney epithelial cells from developing a cystic phenotype.

## DATA SHARING STATEMENT

The datasets are presented in the figures and the supplementary information. The mRNA-seq data are deposited into the Gene Expression Omnibus (GEO) database (GSE262417) and are available online. Additional information is available from the corresponding author on reasonable request.

## ACKNOWLEDGMENTS

The authors would like to thank the patients and their families for their cooperation and interest in the study. This work was supported by grants from NIH/NIDDK (R01DK080745) and a philanthropic gift for PKD research at CCF to OW, Kidney Research UK and the PKD Charity UK (PKD_RP_005_20211124), the Sheffield Hospitals Charity and the Sheffield Kidney Research Foundation to AJS and ACMO, the Deutsche Forschungsgemeinschaft (DFG, BE 3910/8-2, BE 3910/9-1, Project-ID 431984000 - Collaborative Research Center SFB 1453), the Federal Ministry of Education and Research (BMBF, 01GM1903I and 01GM1903G) and the European Union’s Horizon Europe research and innovation programme (grant agreement 101080717, TheRaCil) to CB. DS was supported by a Faculty PhD Scholarship from the University of Sheffield. We thank Drs. S. Somlo, P. Igarashi, and K. Dell for mouse strains, S. Feng, and L. Chang for technical assistance and R. Allen Schweickart for bioinformatical support.

## SUPPLEMENTARY MATERIAL

### Supplementary Figure Legends

**Supplementary Figure S1. *In vitro* Binding Assays Showing Direct Binding between Bicc1, PC1- PLAT and PC1-CT1, but not PC2-CT2.** *In vitro* translated myc-Bicc1 was incubated with recombinant MBP, MBP-PLAT and MBP-CT1 or GST, GST-CT2 and subjected to IP with an anti-c-myc antibody (**a**) or pull-down with GST beads (**b**). MBP or GST was used as a negative control in each respective assay. Arrows indicate pull down of MBP-PLAT and MBP-CT1 respectively; asterisk indicates non-specific band (**a**). GST-CT2 did not bind to myc-Bicc1 directly *in vitro* (**b**). Quality of the different recombinant proteins used is shown by Coomassie staining (**c**). Western blot showing expression of recombinant myc-tagged *Bicc1* generated by *in vitro* translation or myc-tagged *Bicc1* transfected in HEK-293 cells. GST pull-down identified an interaction between co-expressed GST- CT1 and myc-Bicc1 but not with GST (**d**).

**Supplementary Figure S2. Validation of *Xenopus* Knockdowns *and* BICC1 Knockout.** (**a**) qRT-PCR detecting the region targeted by the *Pkhd1-sMO* using a PrimeTime^®^ qPCR assay (IDT). *Xenopus* embryos were injected with the indicated amount of *Pkhd1-sMO* and harvested at stage 39 for mRNA extraction. Individual dots indicate pools of 5 embryos each utilizing three independent fertilizations. Data were analyzed by Mann Whitney test with one asterisk indicating *P* ≤ 0.05 and two asterisks indicating *P* ≤ 0.01. (**b**). To examine cooperativity between Bicc1 and the PKD genes, each MO was titrated for efficacy alone or tested in combination. Embryos were analyzed for edema formation at stage 43. Data are the accumulation of multiple independent fertilizations with the number of embryos analyzed indicated above each condition. Part of the data are shown in **Fig. 3j**. (**c**) qRT-PCR for *BICC1* comparing wildtype cells to the two genetically engineered BICC1 knockout clones. (**d**) qRT-PCR for *PLD2* shows that re-expression of mBicc1, but not the empty vector (pCS2) restored *PKD2* mRNA expression in a HEK293T BICC1 KO clone. (**e,f**) qRT-PCR for *NEFL* and *LAMB3*, which are both downregulated in the HEK293T BICC1 KO clone and restored upon re-expression of mBicc1.

**Supplementary Figure S3. Kidney Parameters of *Bicc1:Pkd2* and *Bicc1:Pkd1* Compound Mutants.** (**a,b**) Comparison of Blood Urea Nitrogen (BUN) levels of kidneys of the *Bicc1:Pkd2* crosses at postnatal day P14 and P21. (**c,d**) Comparison of kidney weight/body weight ratios (KW/BW) levels of kidneys of *Bicc1:Pkd1* crosses and their respective BUN levels at postnatal day P14. (**e**) Immunoprecipitation of PC2 from kidneys of Bicc1^+/+^ and Bicc1^Bpk/Bpk^ mice at postnatal day P4. 200 μg total protein from each sample was used to immunoprecipitate PC2 with 5 μg Ycc2 antibody and agarose-bound protein A/G. PC2 was detected using another antibody against Pkd2 (Sc-28331).

**Supplementary Figure S4. Transcriptomic Analysis of BICC1 Wildtype, BICC1-KO and BICC1- S240P HEK293T cells.** (**a-g**) mRNA-seq data were analyzed using DESeq2 differential expression analysis using two samples per genotype. Venn Diagrams were used to visualize the distribution of the up- or downregulated transcripts (**a, e**); for each intersection the eight most significantly altered transcripts (based on their Padj levels) are visualized in a bar diagram showing the normalized expression levels for each sample (**b-d,f,g**).

**Supplementary Table S1: Expected *vs.* Observed Frequencies in the Bicc1^+/Bpk^:Pkd2^+/+^ x Bicc1^+/Bpk^:Pkd2^+/-^ crosses at P21**.

**Supplementary Table S2: Expected vs. Observed Frequencies in the Bicc1^+/Bpk^:Pkd1^+/+^:Pkhd1- Cre+ x Bicc1^+/Bpk^:Pkd1^+/fl^ crosses at P14**

**Supplementary Table S3: *In Silico* Analysis of the *PKD1* and *PKD2* Variants Identified in VEO- ADPKD Patients.**

**Supplementary Table S4. *In Silico* Analysis of the *BICC1* p.Ser240Pro (S240P) Variant. Supplementary Table S5: Gene Sets Enriched in BICC1-KO *vs.* BICC1-S240P HEK293T Cells.**

**Supplementary Methods**

**Supplementary References**

