## Supplemental Material for "BICC1 Interacts with PKD1 and PKD2 to Drive Cystogenesis in ADPKD"

### SUPPLEMENTARY FIGURES

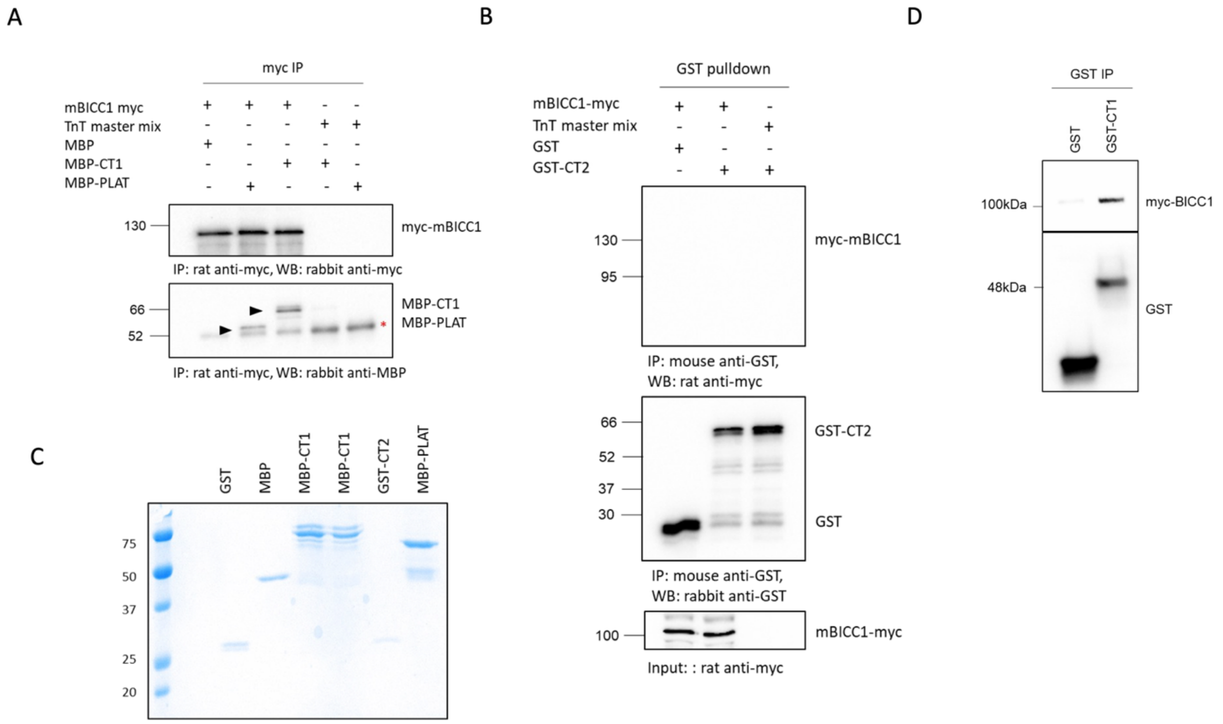


**Supplementary Figure S1. *In vitro* Binding Assays Showing Direct Binding between Bicc1, PC1-PLAT and PC1-CT1, but not PC2-CT2.** *In vitro* translated myc-Bicc1 was incubated with recombinant MBP, MBP-PLAT and MBP-CT1 or GST, GST-CT2 and subjected to IP with an anti-c-myc antibody (**a**) or pull-down with GST beads (**b**). MBP or GST was used as a negative control in each respective assay. Arrows indicate pull down of MBP-PLAT and MBP-CT1 respectively; asterisk indicates non-specific band (**a**). GST-CT2 did not bind to myc-Bicc1 directly *in vitro* (**b**). Quality of the different recombinant proteins used is shown by Coomassie staining (**c**). Western blot showing expression of recombinant myc-tagged *Bicc1* generated by *in vitro* translation or myc-tagged *Bicc1* transfected in HEK-293 cells. GST pull-down identified an interaction between co-expressed GST- CT1 and myc-Bicc1 but not with GST (**d**).

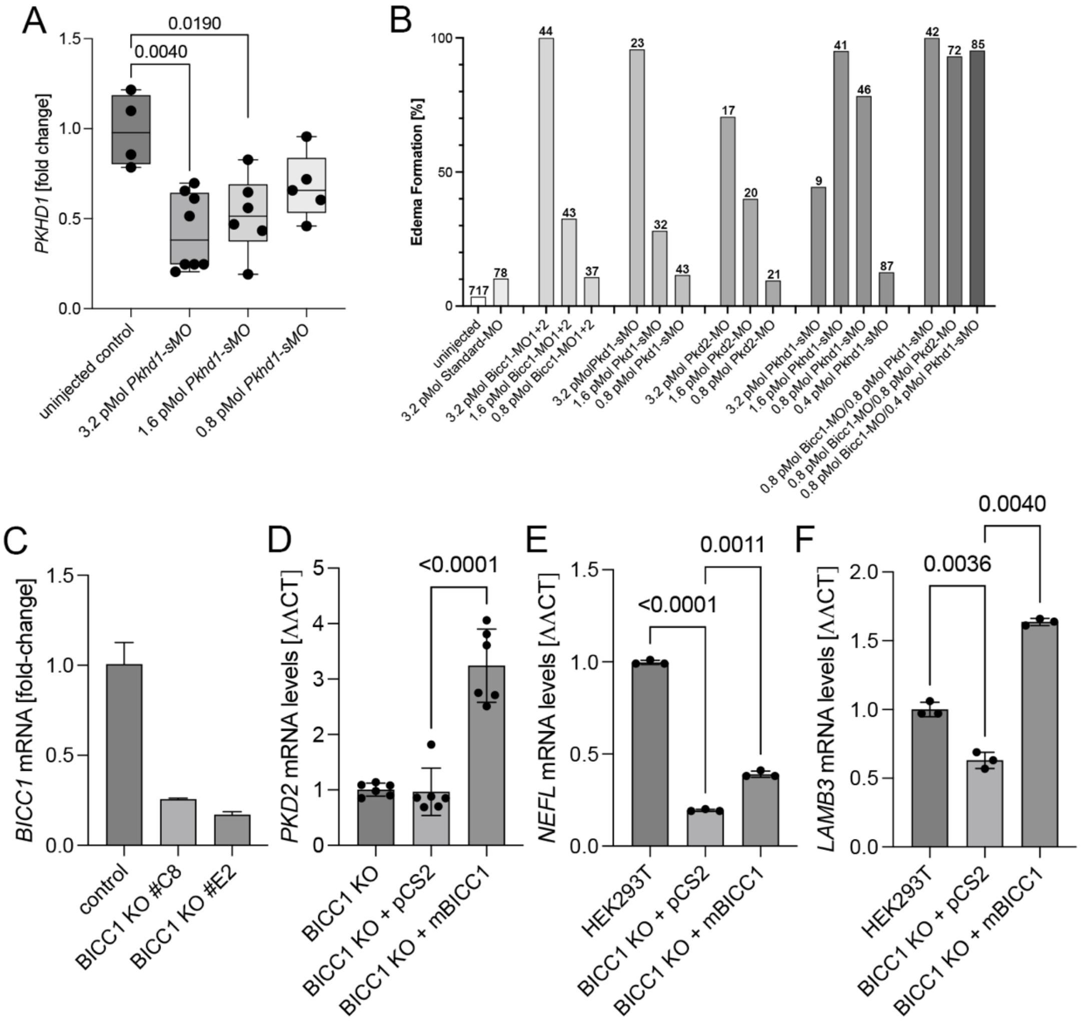


**Supplementary Figure S2. Validation of *Xenopus* Knockdowns *and* BICC1 Knockout.** (**a**) qRT-PCR detecting the region targeted by the *Pkhd1-sMO* using a PrimeTime® qPCR assay (IDT). *Xenopus* embryos were injected with the indicated amount of *Pkhd1-sMO* and harvested at stage 39 for mRNA extraction. Individual dots indicate pools of 5 embryos each utilizing three independent fertilizations. Data were analyzed by Mann Whitney test with one asterisk indicating $P \leq 0.05$ and two asterisks indicating $P \leq 0.01$. (**b**). To examine cooperativity between Bicc1 and the PKD genes, each MO was titrated for efficacy alone or tested in combination. Embryos were analyzed for edema formation at stage 43. Data are the accumulation of multiple independent fertilizations with the number of embryos analyzed indicated above each condition. Part of the data are shown in **Fig. 3j**. (**c**) qRT-PCR for *BICC1* comparing wildtype cells to the two genetically engineered BICC1 knockout clones. (**d**) qRT-PCR for *PLD2* shows that re-expression of mBicc1, but not the empty vector (pCS2) restored *PKD2* mRNA expression in a HEK293T BICC1 KO clone. (**e,f**) qRT-PCR for *NEFL* and *LAMB3*, which are both downregulated in the HEK293T BICC1 KO clone and restored upon re-expression of mBicc1.

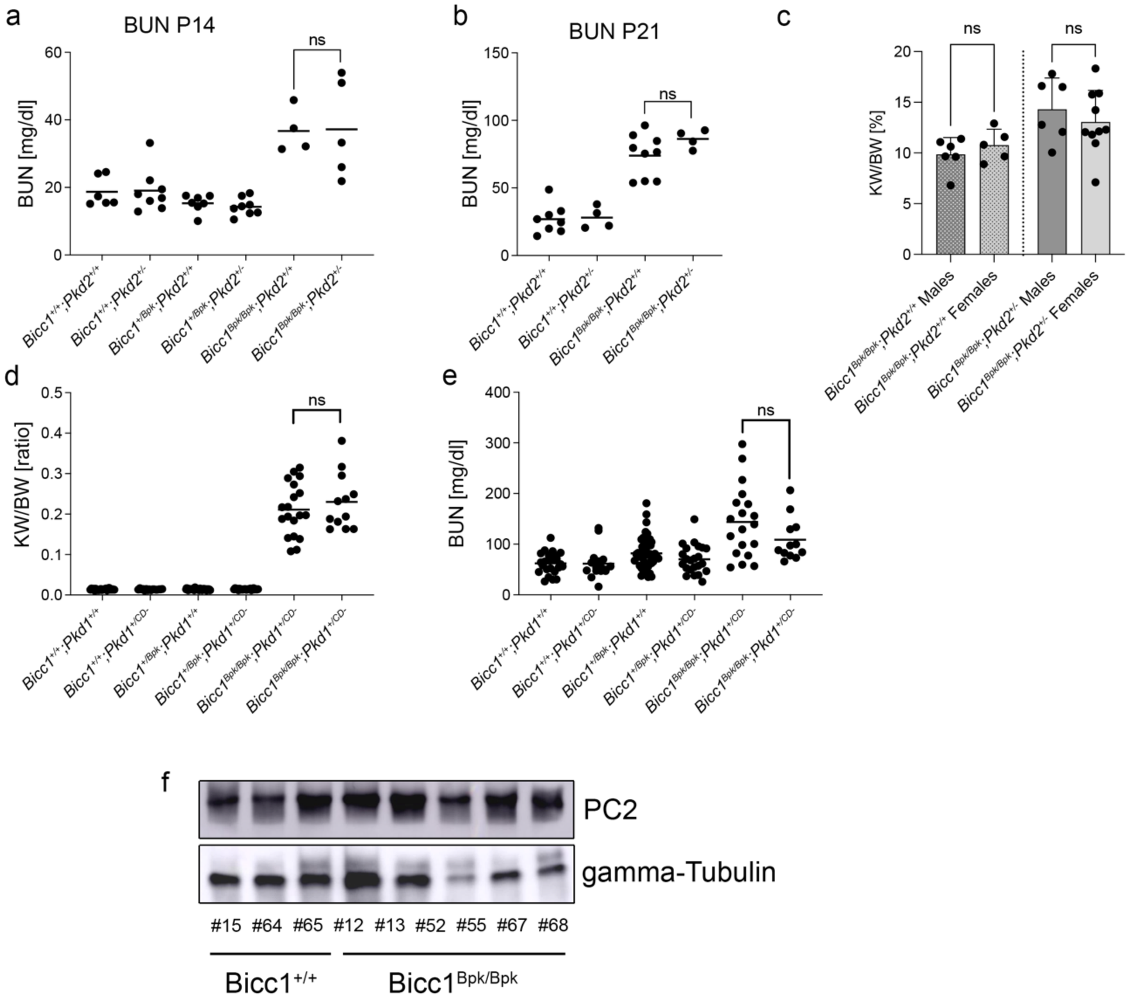


**Supplementary Figure S3. Kidney Parameters of *Bicc1:Pkd2* and *Bicc1:Pkd1* Compound Mutants.** (**a,b**) Comparison of Blood Urea Nitrogen (BUN) levels of kidneys of the *Bicc1:Pkd2* crosses at postnatal day P14 and P21. (**c,d**) Comparison of kidney weight/body weight ratios (KW/BW) levels of kidneys of *Bicc1:Pkd1* crosses and their respective BUN levels at postnatal day P14. (**e**) Immunoprecipitation of PC2 from kidneys of Bicc1$^{+/+}$ and Bicc1$^{Bpk/Bpk}$ mice at postnatal day P4. 200 μg total protein from each sample was used to immunoprecipitate PC2 with 5 μg Ycc2 antibody and agarose-bound protein A/G. PC2 was detected using another antibody against Pkd2 (Sc-28331).

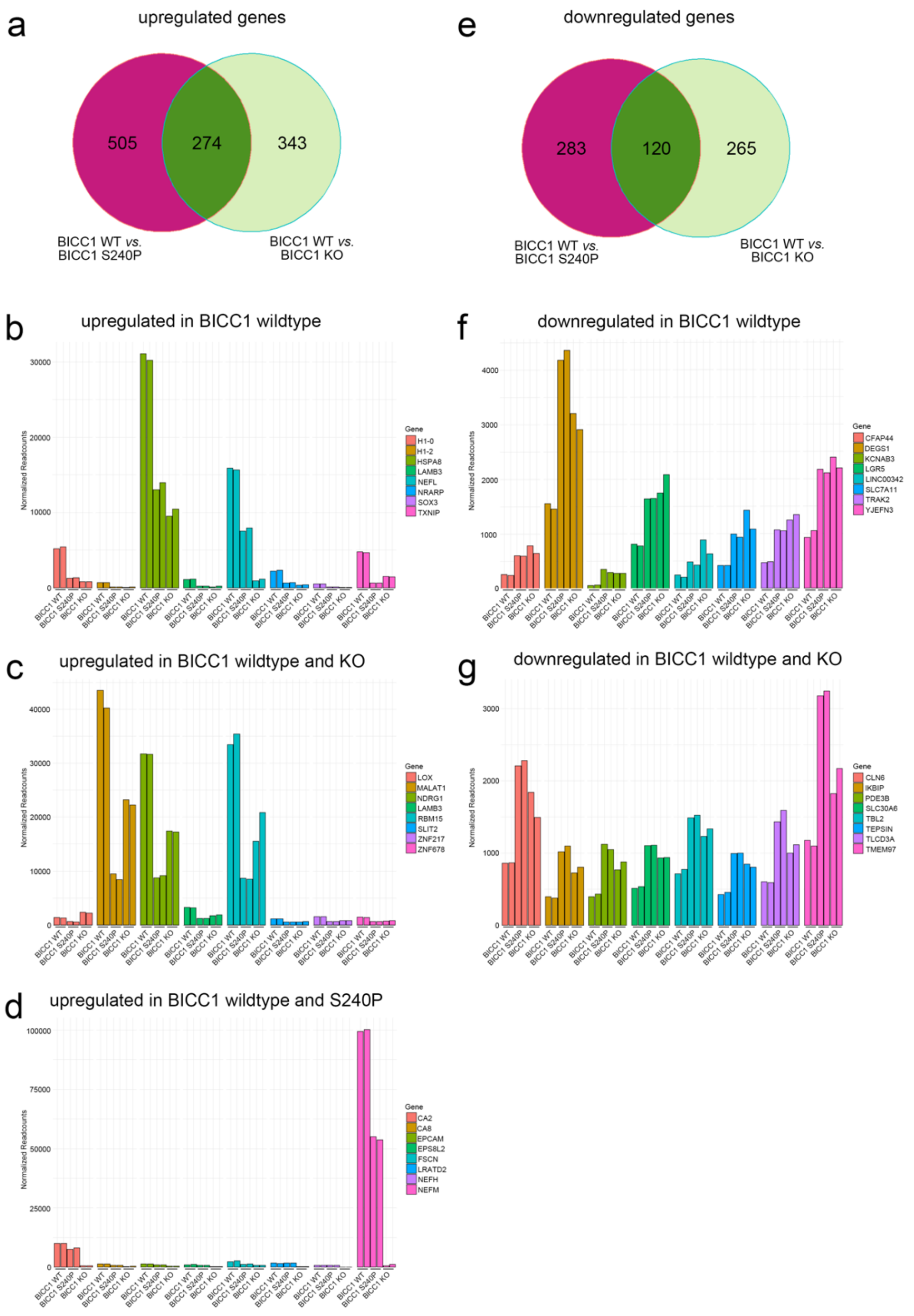


**Supplementary Figure S4. Transcriptomic Analysis of BICC1 Wildtype, BICC1-KO and BICC1-S240P HEK293T cells.** (**a-g**) mRNA-seq data were analyzed using DESeq2 differential expression analysis using two samples per genotype. Venn Diagrams were used to visualize the distribution of the up- or downregulated transcripts (**a, e**); for each intersection the eight most significantly altered transcripts (based on their $P_{adj}$ levels) are visualized in a bar diagram showing the normalized expression levels for each sample (**b-d,f,g**).

### Supplementary Table S1: Expected *vs.* Observed Frequencies in the Bicc1$^{+/Bpk}$:Pkd2$^{+/+}$ x Bicc1$^{+/Bpk}$:Pkd2$^{+/-}$ crosses at P21.

| | Bicc1$^{+/+}$:Pkd2$^{+/+}$ | Bicc1$^{+/+}$:Pkd2$^{+/-}$ | Bicc1$^{+/Bpk}$:Pkd2$^{+/+}$ | Bicc1$^{+/Bpk}$:Pkd2$^{+/-}$ | Bicc1$^{Bpk/Bpk}$:Pkd2$^{+/+}$ | Bicc1$^{Bpk/Bpk}$:Pkd2$^{+/-}$ |
|---|---|---|---|---|---|---|
| Observed [N] | 15 | 8 | 29 | 14 | 12 | 9 |
| Observed [%] | 17.2 | 9.2 | 33.3 | 16.1 | 13.8 | 10.3 |
| Expected [%] | 12.5 | 12.5 | 25.0 | 25.0 | 12.5 | 12.5 |

The observed frequency distribution follows the expected Mendelian distribution; A non-Mendelian distribution is not supported by the Pearson's χ2 test using the Mendel Excel workbook[1] (p=0.159 using a confidence interval of 0.05). Light gray column is the Bicc1$^{Bpk/Bpk}$ phenotype and the dark grey column is the Bicc1$^{Bpk/Bpk}$:Pkd2$^{+/-}$ phenotype. Note that the compound mutants are at a lower than the expected 12.5% frequency.

### Supplementary Table S2: Expected vs. Observed Frequencies in the Bicc1$^{+/Bpk}$:Pkd1$^{+/+}$: Pkhd1-Cre+ x Bicc1$^{+/Bpk}$:Pkd1$^{+/fl}$ crosses at P14

| | Bicc1$^{+/+}$:Pkd1$^{+/+}$: Pkhd1 Cre- | Bicc1$^{+/+}$:Pkd1$^{+/+}$: Pkhd1 Cre+ | Bicc1$^{+/Bpk}$:Pkd1$^{+/+}$: Pkhd1 Cre- | Bicc1$^{+/Bpk}$:Pkd1$^{+/+}$: Pkhd1 Cre + | Bicc1$^{Bpk/Bpk}$:Pkd1$^{+/+}$ : Pkhd1 Cre- | Bicc1$^{Bpk/Bpk}$:Pkd1$^{+/+}$ : Pkhd1 Cre+ | Bicc1$^{+/+}$:Pkd1$^{+/fl}$: Pkhd1 Cre- | Bicc1$^{+/+}$:Pkd1 $^{+/fl}$: Pkhd1 Cre+ | Bicc1$^{+/Bpk}$:Pkd1 $^{+/fl}$: Pkhd1 Cre- | Bicc1$^{+/Bpk}$:Pkd1 +/fl: Pkhd1 Cre+ | Bicc1$^{Bpk/Bpk}$:Pkd1 $^{+/fl}$: Pkhd1 Cre- | Bicc1$^{Bpk/Bpk}$:Pkd1$^{+/fl}$ : Pkhd11 Cre+ |
|---|---|---|---|---|---|---|---|---|---|---|---|---|
| Observed [N] | 8 | 2 | 14 | 17 | 4 | 8 | 11 | 18 | 17 | 23 | 6 | 12 |
| Observed [%] | 5.71 | 1.43 | 10.00 | 12.14 | 2.86 | 5.71 | 7.86 | 12.86 | 12.14 | 16.43 | 4.29 | 8.57 |
| Expected [%] | 6.25 | 6.25 | 12.50 | 12.50 | 6.25 | 6.25 | 6.25 | 6.25 | 12.50 | 12.50 | 6.25 | 6.25 |

The observed frequency distribution does not follow the expected Mendelian distribution; A non-Mendelian distribution is supported by by the Pearson's χ2 test using the Mendel Excel workbook[1] (p=0.019 using a confidence interval of 0.05). Light gray columns are the genotypes yielding a Bicc1$^{Bpk/Bpk}$ phenotype and the dark grey column is the Bicc1$^{Bpk/Bpk}$:Pkd1$^{+/CD}$ phenotype. Note that the mutant phenotype is in line with the expected 6.25% frequency.

**Supplementary Table S3.** ***In Silico Analysis of the PKD1 and PKD2 Variants Identified in VEO-ADPKD Patients.***

| **Gene** | ***PKD2*** | ***PKD1*** | ***PKD2*** |
|---|---|---|---|
| **Chromosomal position** | 4:88046767 | 16:2090945 | 4:88056263 |
| **HGVSc** | c.1445T>G | c.11942C>T | c.1894T>C |
| **HGVSp** | p.Phe482Cys | p.Ala3981Val | p.Cys632Arg |
| **Protein Region** | TRANSMEM-Helical; Range:469-489 | TOPO_DOM-Extracellular; Range:3957-3984 | INTRAMEM-Pore-forming; Range:632-646 |
| **SIFT** | **0.005** | **0.005** | **0.002** |
| **Polyphen2** | 0.519 | **1** | **0.839** |
| **CADD** | **24.9** | **25.9** | **27** |
| **FATHMM** | -0.47 | -0.83 | **-4.45** |
| **Eigen-PC** | 0.450 | 0.356 | **0.538** |
| **GERP++ RS** | **5.61** | 3.12 | **5.36** |
| **EVE** | 0.240 | **0.706** | **0.875** |
| **REVEL** | 0.185 | **0.668** | **0.831** |
| **MetaSVM** | -0.359 | -0.033 | **0.980** |
| **MetaLR** | 0.340 | 0.495 | **0.904** |
| **PrimateAI** | 0.429 | **0.897** | 0.714 |
| **Alphamissense** | 0.088 | **0.503** | **0.993** |
| **ESM1b** | **-7.82** | **-7.96** | **-18.21** |
| **ProtVar** | 1.089 | - | **23.0107** |
| **gnomAD exomes AF** | 0.00204 | 0.00001 | - |
| **gnomAD genomes AF** | 0.00185 | 0.00001 | - |

### Supplementary Table S4. *In silico* Analysis of the *BICC1* p.Ser240Pro (S240P) Variant.

| p.Change (Domain) | Consurf | Amino acid (REF/ALT) | | | | DynaMut ΔΔG | Varsite | |
|---|---|---|---|---|---|---|---|---|
| | | Polarity | Charge | Chemical | HI | | DP | Prediction |
| S240P (KHL1) | 6 | P/NP | N/N | HYDROXYL/ALIPHATIC | -0.8/1.6 | -0.293 | 1.32 | Unfavoured |

NP - nonpolar; P - polar; N – neutral; HI - Hydropathy index; ConSurf - conservation scores (9 - conserved, 1 - variable),; ΔΔG in kcal/mol (change in folding free energy between wild-type and mutant structures, ΔΔG ≥ 0 as stabilizing and ΔΔG < 0 as destabilizing); DP - Disease propensity value (normalized ratio of the number of disease-to-natural variants of a given type).

**Supplementary Table S5: Gene Sets Enriched in BICC1-KO *vs.* BICC1-S240P HEK293T Cells.**

| Rank | Geneset | NOM p-val |
|---|---|---|
| 1 | HALLMARK_EPITHELIAL_MESENCHYMAL_TRANSITION | 0 |
| 2 | HALLMARK_UV_RESPONSE_DN | 0.345 |
| 3 | HALLMARK_ANGIOGENESIS | 0.704 |
| 4 | HALLMARK_KRAS_SIGNALING_DN | 0.357 |
| 5 | HALLMARK_MITOTIC_SPINDLE | 0.345 |
| 6 | HALLMARK_FATTY_ACID_METABOLISM | 0.718 |
| 7 | HALLMARK_IL6_JAK_STAT3_SIGNALING | 0.704 |
| 8 | HALLMARK_APICAL_SURFACE | 0.704 |
| 9 | HALLMARK_TNFA_SIGNALING_VIA_NFKB | 0.704 |
| 10 | HALLMARK_NOTCH_SIGNALING | 0.704 |
| 11 | HALLMARK_INTERFERON_GAMMA_RESPONSE | 0.704 |
| 12 | HALLMARK_INTERFERON_ALPHA_RESPONSE | 0.704 |
| 13 | HALLMARK_P53_PATHWAY | 0.704 |
| 14 | HALLMARK_COAGULATION | 0.704 |
| 15 | HALLMARK_MYOGENESIS | 0.704 |
| 16 | HALLMARK_IL2_STAT5_SIGNALING | 0.704 |
| 17 | HALLMARK_APICAL_JUNCTION | 0.704 |
| 18 | HALLMARK_HYPOXIA | 0.704 |
| 19 | HALLMARK_KRAS_SIGNALING_UP | 0.704 |
| 20 | HALLMARK_ANDROGEN_RESPONSE | 0.704 |
| 21 | HALLMARK_APOPTOSIS | 0.704 |
| 22 | HALLMARK_INFLAMMATORY_RESPONSE | 0.704 |
| 23 | HALLMARK_ESTROGEN_RESPONSE_EARLY | 0.704 |

### Supplementary Methods

#### Cell Culture Studies

UCL93 kidney epithelial cells were immortalized from primary cultures of tubular cells isolated from normal human kidneys removed for clinical indications as previously described.[2,3] Cells were grown in Dulbecco's modified Eagle's medium-Ham's 12 (DMEM-F12, Invitrogen) supplemented with 1% l-glutamine (Invitrogen), 5% NuSerum (Becton Dickinson), and 1% antibiotic/antimycotic solution (Invitrogen) at 33°C/5% $CO_2$.

HEK-293 cells were cultured in Dulbecco's modified Eagle's medium-Ham's 12 (DMEM-F12, Invitrogen) supplemented with 1% l-glutamine (Invitrogen), 10% FCS, and 1% antibiotic/antimycotic solution (Invitrogen) at 37°C/5% $CO_2$. Cells were transfected using Lipofectamine 3000 (Life Technologies) for 48 hours before the cell assays. CRISPR/Cas9 mediated knockout and the BICC1 p.Gly821Glu (BICC1-G821E) and BICC1 p.Ser240Pro (BICC1-S240P) knock-in clonesin HEK293T cells were generated by Synthego Corporation (Redwood City, CA, USA) with the specifics outlined below. The BICC1 knockout was confirmed by qRT-PCR (**Supplementary Figure S2c**) and like in the mouse resulted in a loss of Pkd2 expression that could be rescued by re-expression of mouse Bicc1 (**Supplementary Figure S2d**). In addition, two other genes lost upon elimination of BICC1, *NEFL* and *LAMB3*, were also restored upon re-expression of mouse Bicc1 (**Supplementary Figure S2e,f**). For each engineered cell two independent clones were generated and analyzed. Data were compared to the mock transfected parental cell line. Clonal identity was confirmed at regular intervals using the PCR primers indicated below.

#### Details on Gene Editing of HEK293T cells

#### Bicc1 KO

| | |
|---|---|
| **Cell Line** | HEK293 |
| **Gene Name** | BICC1 |
| **Transcript ID** | ENST00000373886.8 |
| **Guide RNA Sequence** | GAGCGAGGAGCGCUUCCGCG |
| **Guide RNA cut location** | Chr10:58,513,298 |

| | |
|---|---|
| **Exon Targeted** | 1 |
| **PCR & Sequencing Primers** | FOR Primer (5'-3') TGCAGGGGGACGAGCTA<br>REV Primer (5'-3') TGGAGCTAAACCGGCCG |
| **Sequencing Primer** | FOR Primer (5'-3') TGCAGGGGGACGAGCTA |

Genotype Analysis:

1. Clone E1
   Indel: +1
   Description: Homozygous KO clone
2. Clone B8
   Indel: -8/+1
   Description: Compound heterozygous KO clone

**BICC1 carrying p.Ser240Pro (BICC1-S240P)**

| | |
|---|---|
| **Cell Line** | HEK293 |
| **Gene Name** | BICC1 |
| **Transcript ID** | ENST00000373886.8 |
| **Guide RNA Sequence** | UGACAGUAGCACCAUACAUU |
| **Guide RNA cut location** | Chr 10: 58,789,402 |
| **Donor Sequence** | AACCGGTTCCTGATCCTAATTCCCCCTCTATTCAGCA<br>TATATCACAAACGTACAATATTTCAGTACCATTTAAA<br>CAGCGTTCACGAATGTATGGTGCTACTGTCATAGTAC<br>GAGGGTCTCAGAATAACACT |
| **PCR & Sequencing Primers** | FOR Primer (5'-3') TGCTTTAACTCTCTGCTTTGGA<br>REV Primer (5'-3') ACGGGGAAAGATTCTATTGCA |
| **Sequencing Primer** | FOR Primer (5'-3') TGCTTTAACTCTCTGCTTTGGA |

Genotype Analysis:

1. Clone C8
   Modification: BICC1 p.Ser240Pro (TCA>CCA)
   Description: Homozygous KI clone
2. Clone F7
   Modification: BICC1 p.Ser240Pro (TCA>CCA)
   Description: Homozygous KI clone

**BICC1 carrying p.Gly821Glu (BICC1-G821E)**

| | |
|---|---|
| **Cell Line** | HEK293 |
| **Gene Name** | BICC1 |
| **Transcript ID** | ENST00000373886.8 |
| **Guide RNA Sequence** | GACCGAAAUGGAAUUGGACC |
| **Guide RNA cut location** | Chr10:58,813,922 |
| **Donor Sequence** | AGCACTTGGGAGGTGGAAGCGAATCTGATAACTGGAGAG<br>ACCG<br>AAATGAAATTGGGCCTGGAAGTCATAGTGAATTTGCAGCT<br>TCTATT GGCAGCCCTAA |
| **PCR & Sequencing Primers** | FOR Primer (5'-3'): AAAGGCTGTAGGCAGGTTCC |

| | |
|---|---|
| | REV Primer (5'-3'): TCAGAGAGGCCACAGTCAGT |
| **Sequencing Primer** | FOR Primer (5'-3'): AAAGGCTGTAGGCAGGTTCC |

Genotype Analysis:

3. Clone A2
   Modification: BICC1p.Gly821Glu (GGA>GAA)
   Description: Homozygous KI clone
4. Clone E5
   Modification: BICC1 p.Gly821Glu (GGA>GAA)
   Description: Homozygous KI clone

**Transcriptome Analysis**

For mRNA-sequencing, mRNA was extracted using Trizol followed by DNAse treatment. Each cell line/clone was analyzed in triplicates as true technical replicates. Library generation was performed using TruSeq RNA Library Prep Kits (Illumina, San Diego, CA, USA) and sequenced NovaSeq6000 S4 150PE using the services of Psomagen. Primary sequence analysis was performed using Galaxy.[4] Sequence reads were aligned to the human genome (GRCh38) using STAR in Galaxy (Galaxy Version 2.7.10B+galaxy4) with default parameters. Read counts were obtained using FeatureCounts (Galaxy Version 2.0.3+galaxy2) with the default parameters and normalized read count were obtained by DESeq2 (Galaxy Version 2.11.40.8+galaxy0). Gene Set Enrichment Analysis (GSEA) was used to identify normalized enrichment scores of 50 human hallmark gene sets.[5] DESeq2 was used to identify differentially expressed genes (DEGs) and calculate their fold changes (FC), p-values, and false discovery rate (FDR)-adjusted p-values.[6] The sequences data are deposited into the Gene Expression Omnibus (GEO) database (GSE262417) and are available online.

**Plasmids**

Full length PC1 and PC2 plasmids used in this manuscript have been previously reported.[7] Polycystin fusion proteins NT2 (PKD2 aa1–223), NT2 1-100 (PKD2 aa1–100), NT2 101-223 (PKD2 aa101–223), CT2 (PKD2 aa680–968), PLAT (PKD1 aa3118-3223) and CT1 (PKD1 aa4107–4303) were subcloned into pGEX-6P-1, pEBG or pMAL-c2X vectors to express Nterminal bacterial, mammalian GST-fusion proteins, or MBP-fusion proteins respectively.[8,9] myc-mBicc1-ΔSAM (BICC1 aa1-815) and myc-mBicc1-ΔKH (BICC1 aa352-977) truncations were generated by PCR cloning from full-length myc-mBicc1 plasmid. All plasmids were verified by Sanger sequencing. Of note, we have adapted a spelling of Bicc1, where BICC1 is the human homologue, mBicc1 is the mouse homologue and xBicc1 the *Xenopus* one.

**Antibodies**

Primary antibodies used in this study were BICC1 mAb (sc-514846, Santa Cruz), rabbit BICC1 (HPA045212 Sigma), PC1 mAb (7e12),[10] rabbit PC1 (2b7),[11] goat PC2 (sc-10376, Santa Cruz), rabbit PC2 (YCC2, a kind gift from Dr. S. Somlo or Sc-28331, Santa-Cruz Biotechnology), mouse ACTB (ab6276, Abcam), HA High Affinity (rat monoclonal clone 3F10, Roche), mouse GST (sc-138, Santa Cruz), rat c-Myc (clone JAC6, Biorad), rabbit GFP (ab6556, Abcam), mouse V5-tag (clone SV5-Pk1, Biorad) and mouse γ-Tubulin (T6557, Sigma). All primary antibodies were used at 1:1000 unless otherwise stated. Secondary antibodies used in this study include goat anti-mouse IgG (1030-05, Southern Biotech), goat anti-rabbit IgG (4050-01, Southern Biotech), goat anti-rat IgG (3050-01, Southern Biotech) and rabbit anti-goat IgG (P0449, Dako). All secondary antibodies were used at 1:10,000, unless otherwise stated in the results section.

**Protein Biochemistry**

Cells were lysed by extraction at 4°C using the IP lysis Buffer (25 mM NaCl, 150 mM EDTA, 1 mM 0.5% NP40, 1% Triton X-100, pH 7.0) supplemented with a protease inhibitor cocktail (Roche). Immunoblotting and immunoprecipitation were performed as previously described.[11] Biorad ChemiDocTM XRS+ and Image Lab 5.1 software were used for visualization and quantification of proteins of interest. All quantification was carried out on non-saturated bands as determined by the software from 3 independent experiments.

**Recombinant Protein Preparation**

Plasmids were transformed into the E. coli strain BL21-RIPL and recombinant protein expression was induced at 37°C for 3 hours with 0.5 mM IPTG. MBP-tagged, GST fusion and His-tagged proteins were purified with Amylose, Glutathione-Sepharose or Nickel columns, respectively, as previously described.[8]

**Preparation of *in vitro* Translated Bicc1**

Myc-tagged Bicc1 was *in vitro* transcribed and translated with a reticulocyte lysate system TnT SP6 (Promega, USA). Briefly, the plasmid DNA (1 µg) and 50 µl of the reaction mixture were incubated for 90 min at 30°C. Expression of myc-BICC1$_{IVT}$ was determined by Western blotting.

**GST Pull-Down Assays**

One to two micrograms of the bacterial GST fusion protein and 10 µl myc-BICC1$_{IVT}$ were incubated in 300 µl binding buffer (1×TBST with 0.2% Tween20) for 1 hour at RT with gentle rotation. 40 µl of 50% Glutathione Sepharose 4B beads (GE Healthcare) were then added and the mixture was incubated with rotation for an additional hour. The beads were sedimented by centrifugation at 6000 rpm for 2 minutes and washed up to six times with 1 ml volumes of ice-cold PBS. Bound proteins were eluted either using 25 µl of elution buffer or by boiling for 5–10 minutes in reducing sample buffer.

***Xenopus* Embryo Manipulations**

*Xenopus* embryos obtained by *in vitro* fertilization were maintained in 0.1x modified Barth medium[12] and staged according to Nieuwkoop and Faber.[13] *Xenopus* experiments we performed injections using at least three independent clutches per experimental group. Final numbers of animals/experimental group varied as survival was clutch-dependent and animals that did not gastrulate properly or were severely malformed were excluded from subsequent analysis. Microinjections were performed on randomly selecting cleaving embryos at the 2- to 4-

cell stage for a given antisense MO/MO combination. Data analysis was performed in a blinded fashion and groups were only revealed post data acquisition. The sequences of the antisense morpholino oligomers (GeneTools, LLC) used in this study were 5'-GGG ACA AAG ATG CTC ATT TTA ACA G-3' (*BicC-MO1*)[14], 5'-GCC ACT ATC TCT TCA ATC ATC TCC G-3' (*BicC-MO2*)[14], 5'-TCC TTA TGG TCC GAG TTA CCT TGG G-3' (*Pkd1-sMO*)[9,15], 5'- GGT TTG ATT CTG CTG GGA TTC ATC G-3' (*Pkd2-MO*)[16] and 5'- TAT TGT GTT CTA TTC TTA CCT TTC T-3' (*Pkhd1-sMO*). For complete knockdown a total of 3.2 pMol of *Std-MO*, *Pkd1-sMO*, *Pkd2-MO*, *Pkhd1-sMO* or a mixture of 3.2 pMol *Bic-C-MO1* and 3.2 pMol *Bic-C-MO2* (*Bic-C-MO1+2*) was injected radially at the 2- to 4-cell stage into *Xenopus* embryos. Note that *Xenopus laevis* is allotetraploid, and while we normally target both the L and S allele with one MO, in the case of Bicc1 it requires two. For suboptimal knockdowns 0.8 pMol of the *Bic-C-MO1*, *Bic-C-MO2*, *Pkd1-sMO* or *Pkd2-MO* and 0.4 pMol *Pkhd1-sMO* were used. Knockdown of Pkd1 and Pkhd1 was performed using MOs targeting 3' splice donor sites (*Pkd1-sMO* and *Pkhd1-sMO*). Microinjection assays and RT-PCR demonstrated that both splice MOs are functional and prevent proper splicing of the two genes (**Supplementary Figure S2a** and Supplementary Figure S12 in [9]). Suboptimal concentrations were determined by injecting serially diluted MOs and determining the concentration-dependent induction of the edema phenotype (**Supplementary Figure S2b**).

For synthetic mRNA, *pCS2-xBicC*[*][14] and its derivatives carrying the corresponding point mutations (generated by Quikchange II Mutagenesis kit from Stratagene) were linearized with *Not*I and transcribed with SP6 RNA polymerase using the mMessage mMachine® (Ambion). Rescue experiments, whole mount in situ hybridizations and histology were performed as previously described.[14] To generate antisense probes the plasmids were linearized and transcribed as follows: *pSK-Bicc1* [17] *- NotI/T7, pCMV-SPORT6-Nbc1* [18] - *Sal*I/T7, *pGEM-T-Easy-Pkd1 - Nco*I/Sp6, *pCRII-TOPO®-Pkd2* [16] - *Not*I/Sp6, *pGEM-T-Easy-Pkhd1 - Nco*I/Sp6.

**Mouse Studies**

For the mouse studies, the sample sizes for the experimental groups were not determined *a priori* as we did not know the effect sizes for the phenotypes under investigation. Thus, we collected multiple litters until the number of the mutant phenotypes were statistically significantly different from the controls and the number of animals in the experimental groups of interest exceeded 10. Genotyping was performed after collecting the biological data; thus, the investigator was blinded during the data acquisition phase. No outliers were removed unless mice were moribund before sacrifice. The Pkd2/Bicc1 mouse crosses were performed using two mouse strains, one carrying the Bicc1 hypomorphic allele Bpk[19] and one of a Pkd2 null allele.[20] As the two mice strains were of different genetic background (BALB/c and C57BL/6, respectively) we utilized a breeding scheme minimizing the influence of the genetic background. *Bicc1*$^{+/Bpk}$ and *Pkd2*$^{+/-}$ mice were crossed to generate *Bicc1*$^{+/Bpk}$*:Pkd2*$^{+/-}$ compound heterozygotes as F1 generation. These mice were then intercrossed to generate the experimental animals in the F2 generation. Mice were genotyped by PCR and analyzed at postnatal day P4, P14 and P21. Kidneys were examined as previously described[16] for kidney function using BUN (QuantiChrom™ Urea Assay Kit, BioAssay Systems), morphometric parameters (body and kidney weight) as well as histology and immunofluorescence analyses (i.e. *Lotus tetragonolobus* agglutinin [LTA] and *Dolichos biflorus* agglutinin [DBA] to determine cyst origin). Cystic index was calculated as percent of the kidney occupied by proximal (LTA-positive) or collecting duct (DBA-positive) cysts.

The *Pkd1/Bicc1* mouse crosses were performed using the same *Bicc1* hypomorphic allele *Bpk*, which was transferred into the C57BL/6 background by backcrossing for more than 10 generations. The *Bpk* allele displayed the same cystic kidney phenotype in this background as the one described for BALB/c.[21] These mice were intercrossed to the *Pkd1*$^{fl/fl}$*;Pkhd1-Cre* mice (a kind gift from Drs. Somlo and Igarashi), an allele we refer to as *Pkd1*$^{CD-}$ in this study. Kidneys were analyzed at postnatal day P7 and P14 for kidney function, morphometric parameters, histology, and immunofluorescence, as described for the *Bicc1/Pkd2* mutants.

Of note, the choice of the mouse strains was based on the availability of mice at the time of the experiments and not due to scientific reasons. As we had not finished backcrossing the Bicc1-Bpk strain from Balb/c into C57BL/6, it would have been scientifically unsound to assume genetic homogeneity and cross them with the Pkd2 mutant mice in an uncontrollable fashion. Thus, the interaction between Bicc1 and Pkd2 was performed by generating breeders ($Bicc1^{+/Bpk}$:$Pkd2^{+/+}$ and $Bicc1^{+/Bpk}$:$Pkd2^{+/-}$) in the F1 generation and the experimental animals in the F2 generation. Yet, when we started exploring the interaction between Bicc1 and Pkd1, all three mouse strains ($Bicc1^{+/Bpk}$, $Pkd1^{fl/fl}$ and Pkhd1-Cre) were available in the C57BL/6 strain and the $Bicc1^{+/Bpk}$ had been backcrossed into C57BL/6 more than 10 generations. Thus, the Bicc1-Pkd1 study was performed using traditional breeding schemes.

**International Diagnostic Clinical Cohort**

Next Generation Sequencing (NGS) technologies and comprehensive bioinformatic analyses utilized in this project are described in detail elsewhere.[22,23] In brief, we performed different NGS-based approaches utilizing a customized sequence capture library with curated target regions - currently comprising of more than 650 genes described and associated with cystic kidney disease or allied disorders - as well as corresponding flanking intronic sequence according to the manufacturer´s recommendations. The panel design is enriched by targets in non-coding regions for described variants listed in well-accepted databases like HGMD or ClinVar and optimized for low-performance and disease-critical regions (e.g., *PKD1*). DNA samples were enriched using sequence capture, multiplexed, and in most cases sequenced using Illumina sequencing-by-synthesis technology with an average coverage of more than 300X. Raw data were processed following bioinformatics best practices. Mapping and coverage statistics were generated from the mapping output files using standard bioinformatics tools (e.g.

Picard). Statistical analysis was conducted on our internal database currently comprising > 20,000 datasets. The total of this data pool is summarized over samples analysed by NGS-based customized panel testing or whole exome sequencing (WES) analysis. Customized panel setups have been regularly updated. Sub-cohorts of patients were categorized based on clinical, ultrasound and/or histologic data. Control cohorts were selected by ruling out any involvement of kidney related symptoms. This approach yielded high and reproducible coverage enabling copy number variation (CNV) analysis. Performance of the wet-lab and bioinformatic processes are validated and controlled according to national and international guidelines[55,56] reaching high sensitivity for SNV, Indels and CNVs using well-established reference samples, as well as a large cohort of positive controls, especially for CNVs.[24,25] For interpretation of identified variants, we established a bioinformatic algorithm automatically calculating ACMG classification based on existing and updated guidelines[26,27] and was conducted according to specific standardized internal procedures. Bioinformatically called variants were classified according to ACMG/AMP and ACGS guidelines in respect to current literature and database entries (internal and external mutation and frequency databases, public clinical and functional studies) as well as family history and - if available - segregation results. Variant prioritization was based on this classification and on the frequency of the respective variants in public databases. Variants (e. g., in the genes *PKD1, PKD2* and *BICC1* were filtered and prioritized for very rare variants in external (gnomAD) and internal databases in our cohort of patients with PKD, classified as pathogenic, likely pathogenic or VUS, not present in the overall control cohort of all patients in our database and/or patients not affected by PKD or a similar phenotype. Sequence variants of interest were verified by Sanger sequencing, if NGS results failed internal validation guidelines.

For statistical analyses of our patient data, we screened our entire internal database. In a control sub-cohort rigorously screened against any clinical involvement of kidney symptoms (>

10,700 patients) neither a *BICC1* variant (Class III-V) in combination with a *PKD1* or *PKD2* variant nor a relevant monoallelic *BICC1* variant could be identified using the workflow used for variant prioritization described above. We also repeated both queries on cohorts of patients clinically presenting as glomerular disease/focal segmental glomerular sclerosis (FSGS) or atypical hemolytic uremic syndrome (aHUS) with 957 and 1,889 cases and datasets, respectively. Again, we did not detect a single patient with any of the variants described in the manuscript.

#### ***In silico*** **Studies**

The 3D structure of BICC1 (UniProt: Q9H694), PKD1 (UniProt: P98161) and PKD2 (UniProt: Q13563) was downloaded from PDB (6GY4, 4RQN, Bicaudal-C ortholog GLD-3 “3N89”, 6A70 and 6WB8), modeled by AlphaFold and the PHYRE2 automated protein homology modeling server.[28-31] Because no experimentally mutant BICC1 structures have been determined, we generated mutant structures by individually introducing the missense mutations *in silico*; missense mutations were then computationally modeled in UCSF Chimera 1.14 [32] by first swapping amino acids using optimal configurations in the Dunbrack rotamer library[33] and by taking into account for the most probable rotameric conformation of the mutant residue. All kinds of direct interactions, i.e. polar and nonpolar, favorable and unfavorable, including clashes, were analyzed using the contacts command in UCSF Chimera 1.14.[32] The evolutionary conservation score of each amino acid of BICC1 in its conserved domains (KH, KHL and SAM domains) was determined using the ConSurf algorithm, based on the phylogenetic relationships between sequence homologues.[34] To determine the effects of the mutations in flexible conformations of the protein, we used DynaMut, a consensus predictor of protein stability based on the vibrational entropy changes predicted by an elastic network contact model (ENCoM).[35] Pathogenicity of the variants was predicted using Ensembl Variant Effect Predictor (VEP)[36] to calculate a REVEL score[37] and the structural impact of missense variants analyzed using

VarSite.[38] The pathogenicity score of BICC1, PKD1 and PKD2 variants was also determined using different predictors with the scores collated from Argus dbNSFP and ProtVar.[39,40]
